# Ploidy-dependent modulation of DNA replication kinetics in *Xenopus*

**DOI:** 10.64898/2026.01.10.698760

**Authors:** Hemalatha Narrissamprakash, Olivier Haccard, Zhen Li, Nicolas Pollet, Kathrin Marheineke

## Abstract

DNA replication in multicellular organisms follows a tightly regulated spatio-temporal program. Although the mechanisms underlying this replication program remain only partially understood, studies in model systems such as *Xenopus laevis* have highlighted the importance of titrating low-abundance replication factors. Whole-genome duplications as a consequence of polyploidization introduce additional layers of complexity, yet comparative analyses of replication programs across closely related polyploid species are scarce. Here, we developed an interspecies *in vitro* replication system using *X. laevis* egg extracts and sperm nuclei from three *Xenopus* species with varying ploidy: diploid (*X. tropicalis*), tetraploid (*X. laevis*), and dodecaploid (*X. eysoole*). Replication in diploid *X. tropicalis* was faster than in tetraploid *X. laevis* nuclei, due to higher fork density and speed. Surprisingly, dodecaploid *X. eysoole* replication was also accelerated compared to tetraploid nuclei, suggesting that higher ploidy does not necessarily extend S phase. Since replication genes are highly conserved between these species, these results imply dynamic tuning of replication programs across polyploid species and shed light on the evolutionary adaptability of DNA replication in response to genome duplication.

## Introduction

DNA replication in multicellular organisms initiates from thousands of sites known as replication origins. Their activation follows a highly conserved pathway that has been characterized in great molecular detail in model systems such as budding yeast, *Drosophila*, *Xenopus laevis* frogs, and mammalian cell cultures. The coordination of more than fifty different protein factors is necessary to (i) license, (ii) activate (fire) a replication origin, (iii) establish two replication forks, and (iv) achieve the faithful duplication of the genetic material (for review see [1]). In late mitosis and G1 phase, origins are licensed for replication by loading the pre-replicative complex (pre-RC) onto chromatin. The pre-RC is composed of the six ORC (origin recognition complex) subunits, Cdc6 (cell-division-cycle 6), and the MCM (mini-chromosome maintenance) 2–7 helicase complex. In metazoans, Donson, Mtbp (Mdm2-binding protein)/Treslin (Ticrr), and TopBP1 are loaded onto the pre-RC to build the pre-initiation (pre-IC) complex and facilitate recruitment of GINS-Cdc45 to form the active replicative Cdc45/MCM2-7/GINS (CMG) helicase complex. The action of cyclin- and Dbf4/Drf1-dependent kinases (CDKs and DDKs) is necessary for the assembly of this complex and its activation. The association of RecQL4 [2] and MCM10 initiates DNA synthesis by recruiting DNA polymerases and elongation factors. The replication timing regulator Rap1-interacting factor (Rif1) has been shown to inhibit late replication origins by recruiting protein phosphatase 1 (PP1) to the vicinity of origins, thereby counteracting the DDK-dependent activation of the replicative helicase MCM2-7 complex [3–5]. In eukaryotes, replication follows a strictly regulated spatio-temporal program that involves the coordinated activation of groups of origins or clusters, as well as larger domains in a Rif1-dependent manner.

For more than three decades, DNA replication has been extensively studied using the *X. laevis* egg extract system [6]. This powerful *in vitro* model accurately reproduces early embryonic cell cycles characterized by very short S phases. In this system, isolated *X. laevis* sperm nuclei replicate synchronously in interphase S-phase extracts obtained from *X. laevis* eggs. Typically, origins are not strictly defined by specific DNA sequences [7] and are spaced 10–15 kb apart [8–10], whereas in somatic mammalian cells, this distance is about tenfold larger [11], consistent with 20–30 times longer S phases. Manipulating this system has provided key insights into the control of S-phase duration and replication dynamics. Increasing the concentration of sperm nuclei in the egg extracts prolongs the S phase [12]. We demonstrated that this is due to a change in the temporal replication program (asynchrony of cluster activation) and, to a lesser extent, the spatial program (inter-origin spacing within clusters) [13]. During *X. laevis* embryogenesis, the S-phase lengthens after the first 12 cell cycles, coinciding with zygotic genome transcription starting at the mid-blastula transition (MBT). Around the MBT, following an increase in the nuclear-to-cytoplasmic ratio, the levels of Treslin, RecQL4, TopBP1, and the embryonic DDK regulatory subunit Drf1 become limiting for replication, resulting in a longer S-phase [14]. DNA fiber analysis of sperm nuclei replicated in *X. laevis* embryonic extracts confirmed that shortly after the MBT, replication origin density decreases by one-third while inter-origin distances increase [15]. Together with numerical simulations in the *in vitro* system [16,17], these findings demonstrate that diffusible, rate-limiting maternal replication factors regulate S-phase duration and the replication program during early development, when transcription has not yet begun. Thus, both the spatial and temporal replication programs change during embryogenesis and cell differentiation in multicellular organisms.

Polyploidization represents a major evolutionary mechanism that reshapes genome structure and drives speciation across eukaryotes [18,19]. While most studies in animal polyploidy have focused on triploids and tetraploids, higher-order polyploidy is frequent among amphibians [20]. Frogs of the genus *Xenopus* are particularly suitable for investigating how genome duplication influences cellular processes, as they encompass species with ploidy levels ranging from diploid (2n=2x) to dodecaploid (2n=12x). The most recent vertebrate whole-genome duplication occurred within this lineage [21–23]. The diploid *X. tropicalis* (2n=2x=20 chromosomes) and the allotetraploid *X. laevis* (2n=4x=36 chromosomes) are well-established model systems. In contrast, the dodecaploid species *X. eysoole* (2n=12x=108 chromosomes) was described more recently [21]. In animals, the impact of chromosomal DNA content on S-phase duration has been analyzed mainly in highly polyploid somatic cells (100–1000N) undergoing endoreduplication (G/S cycles without mitosis), such as in specialized cells of the fruit fly *Drosophila* and mice [24]. This study and others (reviewed in [25]) revealed that the S phase can either slow down (in *Drosophila*) or remain unchanged (in mice). Interestingly, early embryonic cell cycles in *X. tropicalis* and *X. laevis* are similar in length [26], yet the replication programs of *Xenopus* species with different ploidy have never been compared in detail.

Here, we established an interspecies *in vitro* replication system using *X. laevis* interphase egg extracts and sperm nuclei from three *Xenopus* species representing different ploidy levels: diploid (*X. tropicalis*), tetraploid (*X. laevis*), and dodecaploid (*X. eysoole*). We characterized their replication kinetics, origin activation, and fork progression. We found that *X. tropicalis* sperm nuclei replicate faster than those of *X. laevis*, owing to an increased rate of the replication program, a global increase in the number of activated replication origins, and higher fork speeds. Surprisingly, replication of *X. eysoole* nuclei was also accelerated relative to the tetraploid species, and similarly to the diploid species, indicating that higher ploidy does not necessarily slow down replication kinetics and prolong S phase. A comparative analysis of the conservation and retention of replication-related genes across the three *Xenopus* species suggests that dosage-compensation mechanisms have evolved to adapt DNA replication to different polyploidy levels.

## Results

### Interspecies *in vitro* system to study the effect of different polyploid levels in *Xenopus tropicalis* (2x), *Xenopus laevis* (4x), and *Xenopus eysoole* (12x): Genome size-nuclear size correlation

Previous work has demonstrated that increasing the concentration *of X. laevis* sperm nuclei in egg extracts slowed down the replication program, suggesting that replication factors become rate-limiting [13,27]. We now wanted to study the effects of increasing genome size on DNA replication using three closely related *Xenopus* species. We isolated sperm nuclei from the diploid *X. tropicalis* (3.6 pg DNA/nucleus), the allotetraploid *X. laevis* (6.35 pg DNA/nucleus), and the dodecaploid *Xenopus eysoole* (16 pg DNA/nucleus) (Fig. 1A). We measured the mean nuclear area of Hoechst-stained isolated sperm after fluorescence microscopy (Fig. 1B). As described before for other nuclei types of different *Xenopus* species [28], nuclear size increased significantly and linearly between each species with increasing polyploidy and chromosome content (Fig.1 C, D), showing a clear correlation between genome size and nuclear scaling.

**Fig. 1:**
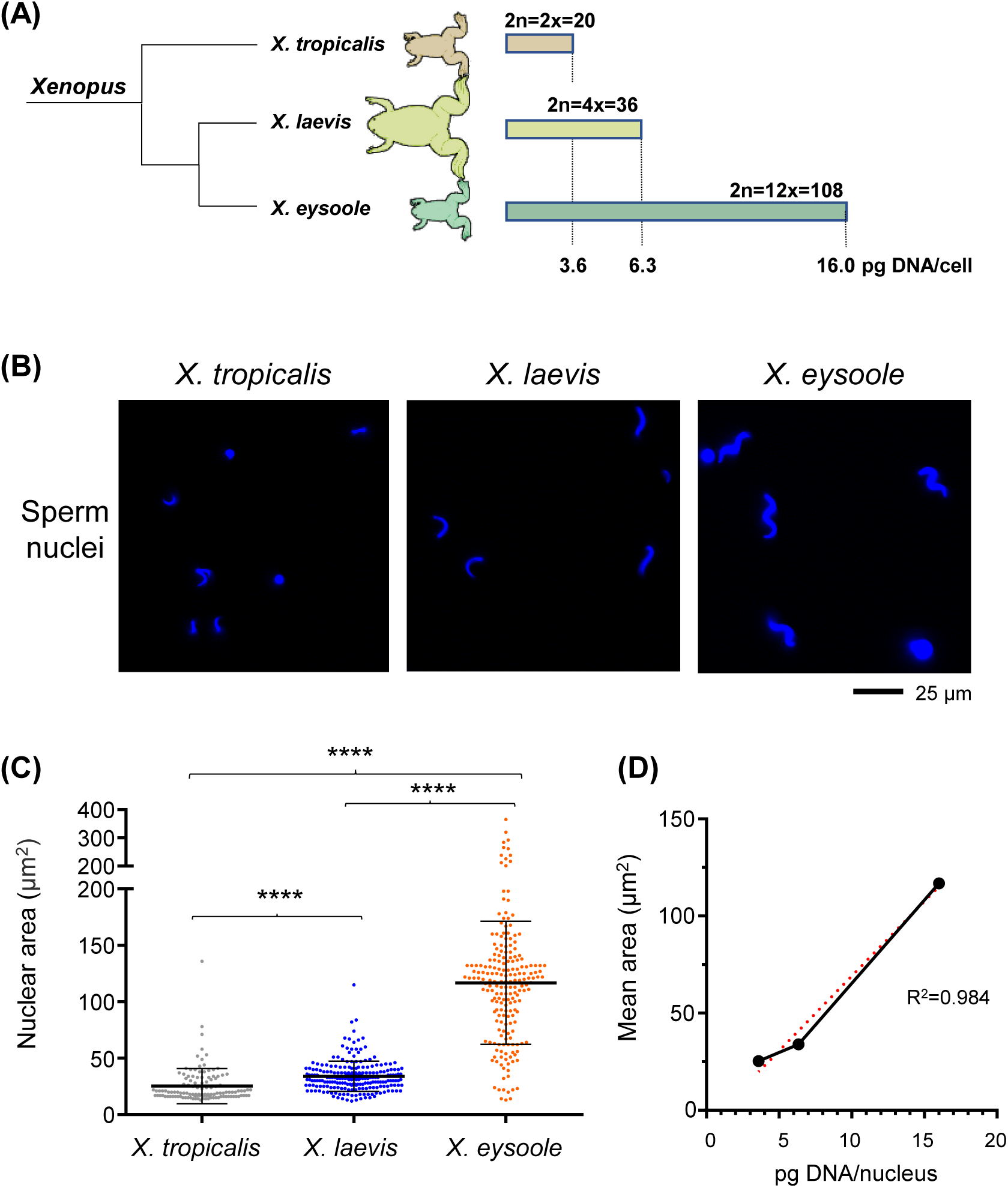
Linear relationship between genome size and nuclei size in *X. tropicalis*, *X. laevis*, and *X. eysoole* isolated sperm. (A) Phylogeny of *X. tropicalis, X. laevis*, and *X. eysoole* with indicated chromosome number and DNA content. (B) Representative fluorescence-microscopy images of isolated sperm nuclei from the three species, Hoechst-stained. (C) Quantification of sperm nuclear area (µm²) shown as a scatter plot with mean ± SD; n = 121, 226, 204; scale bar = [25 µm]. (D) Average nuclear area from C as a function of DNA content, fitted with linear regression in red and displayed with R² value; Y = 7,696 *X - 7,885. An asterisk indicates significant difference (Mann-Whitney U test, two-sided, p<0.05): p-values: * 0.01-0.05; ** 0.001-0.01; *** 0.0001-0.001; **** < 0.0001).

### Faster replication kinetics of the diploid *X. tropicalis* compared to tetraploid *X. laevis* sperm nuclei in *X. laevis* egg extract

To compare replication kinetics, we first focused on *X. laevis* (4x) and *X. tropicalis* (2x) nuclei. Sperm nuclei from both species at the same concentration (2000 nuclei/µl) were incubated in S-phase egg extract in the presence of α[³²P]-dCTP, reactions were stopped at different times during S phase (Fig. 2A), and DNA synthesis was quantified after DNA-gel electrophoresis. Upon the addition of permeabilized sperm DNA to egg extracts, chromatin and replication proteins are imported into the nucleus, chromatin is assembled, and replication proteins are recruited to the chromatin; finally, nuclei synchronously start DNA replication. Replication kinetics experiments revealed that nuclei from both species entered S phase at similar times, but the slope of α[³²P]-dCTP-incorporation was steeper for diploid *X. tropicalis* nuclei than for tetraploid *X. laevis* nuclei, indicating that DNA synthesis progressed more rapidly in *X. tropicalis* (Fig. 2B, C).

**Fig. 2:**
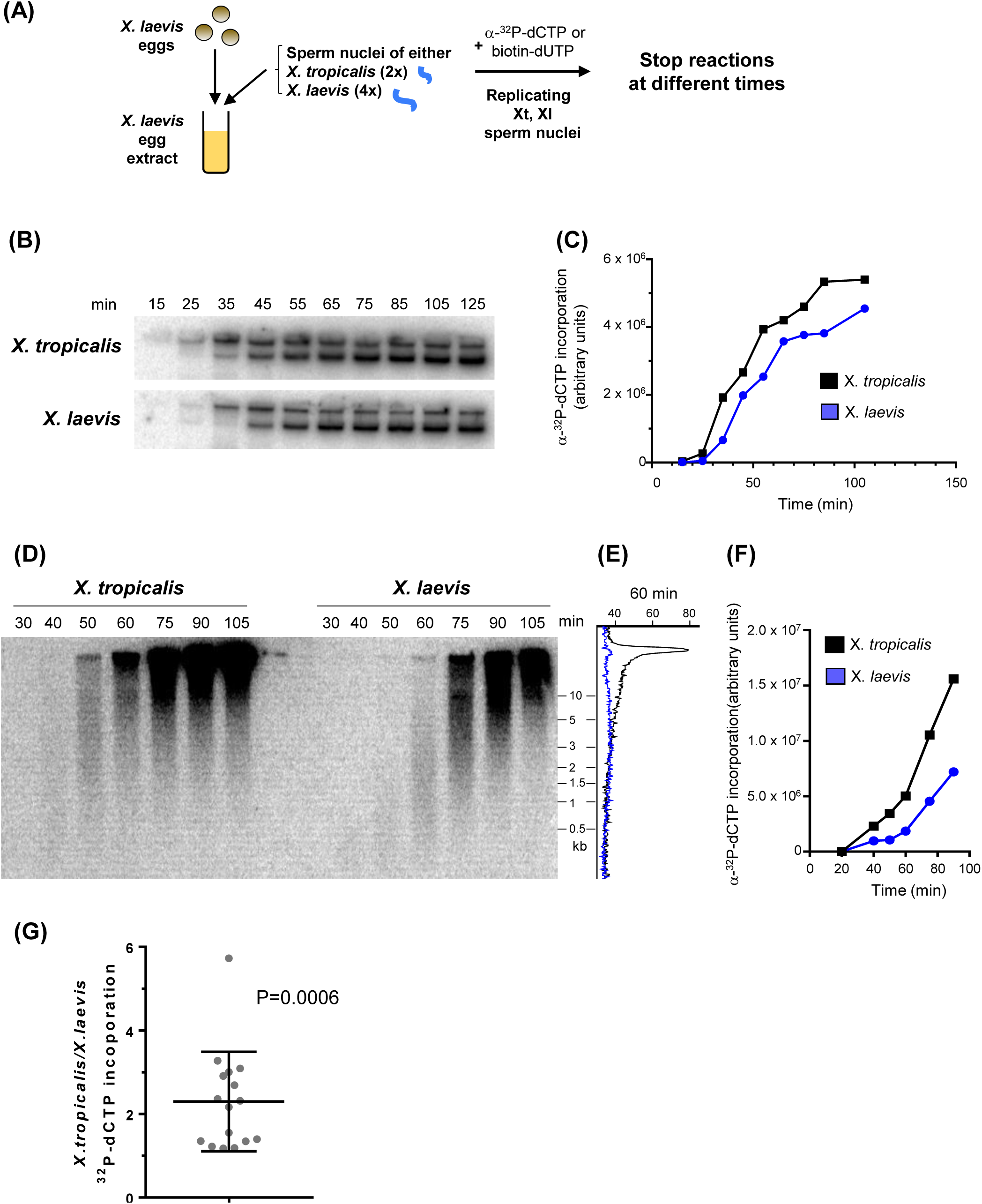
Higher replication rate in *X. tropicalis* sperm than in *X. laevis* sperm in *X. laevis* egg extract. (A) Sperm nuclei (2000 nuclei/µL) from *X. tropicalis* and *X. laevis* were incubated in *X. laevis* egg extracts supplemented with α[³²P]-dCTP. Replication reactions were stopped at the indicated time. (B) Genomic DNA was purified and separated by agarose gel electrophoresis. (C) Quantification of incorporated α[³²P]-dCTP (arbitrary units) in (B). (D) Same as (B), but replicated, genomic DNA from *X. tropicalis* and *X. laevis* was separated on an alkaline gel. (E) Greyscales line profiles of 60 min lanes of *X. laevis* (blue) and *X. tropicalis* (black) from (D), strand sizes indicated in kb. (F) Quantification of α[³²P]-dCTP (arbitrary units) in (D). (G) α[³²P]-incorporation *X. tropicalis/X. laevis* ratios from (C) and (F), scatter plot with mean and SD, one-sample t-test, p-value indicated; n=16.

To visualize nascent strand DNA synthesis, we performed alkaline gel electrophoresis, focusing on early to mid S phase in a second independent α[³²P]-dCTP-incorporation experiment (Fig. 2D-F). We found that nuclei from both species started DNA synthesis at similar times, confirming that nuclear assembly and import of nuclear factors occurred at comparable rates. However, nascent strands from *X. tropicalis* reached higher molecular weights more quickly than those from *X. laevis* (Fig. 2E). Replication rate was higher during early and mid S phase (Fig. 2F). On average, a significant 2.3-fold increase of DNA synthesis was observed in *X. tropicalis* compared to *X. laevis* nuclei (Fig. 2G). This increase could be attributed to either a faster fork speed or a higher initiation rate, leading to quicker merging of neighboring replication forks.

To distinguish between these possibilities, we performed DNA combing experiments in the same extract, parallel to the α[³²P]-dCTP-incorporation experiment shown in Fig. 2D-F. DNA combing analysis allows visualization of origin activation and fork progression on single DNA fibers (Fig. 3A). Sperm nuclei from *X. tropicalis* and *X. laevis* were incubated in S-phase egg extracts in the presence of biotin-dUTP, reactions were stopped at different times, and DNA fibers were immunostained and analyzed (Suppl. Table 1). As seen above, the replication extent in *X. tropicalis* was higher than that of *X. laevis* sperm (Fig. 3B, Suppl. Table 1). Quantification of the number of active replication forks revealed 3.7-fold more active forks in *X. tropicalis* than in *X. laevis* sperm in mid-S phase (Fig. 3C), indicating that more origins were globally active throughout the genome. Comparison of eye-to-eye distances, which serves as a local measure of origin distances by DNA combing, showed that no significant difference was detectable between the two species (Fig. 3D). Since more replication forks were detected in *X. tropicalis* at that time point, these results suggest that the activation of replication clusters occurred more synchronous in *X. tropicalis*, potentially shortening the temporal replication program compared to *X. laevis*. Eye lengths were significantly larger in *X. tropicalis* than in *X. laevis* (Fig. 3E), indicating either a faster fork speed or quicker merging of neighboring replication eyes in *X. tropicalis*. Next, we compared replication parameters at equal fractions of replicated DNA fibers between the two species to determine whether they change intrinsically at the level of replication clusters (Fig. 3F) [29]. This analysis showed that the average fork density within DNA fibers, or inside replication clusters, peaks at lower replication extents and is lower at high replication extents for *X. tropicalis* than for *X. laevis* (Fig. 3G). Additionally, eye-to-eye distances are slightly smaller for *X. laevis* than for *X. tropicalis* at higher replication extents (Fig. 3H). These results show that in *X. tropicalis,* a lower number of activated origins at larger replication extents inside replication clusters partially counterbalances the general acceleration of the temporal program. In conclusion, replication kinetics are faster in diploid *X. tropicalis* sperm than in *X. laevis* sperm due to a higher global initiation rate and an increase in the apparent fork speed.

**Fig. 3:**
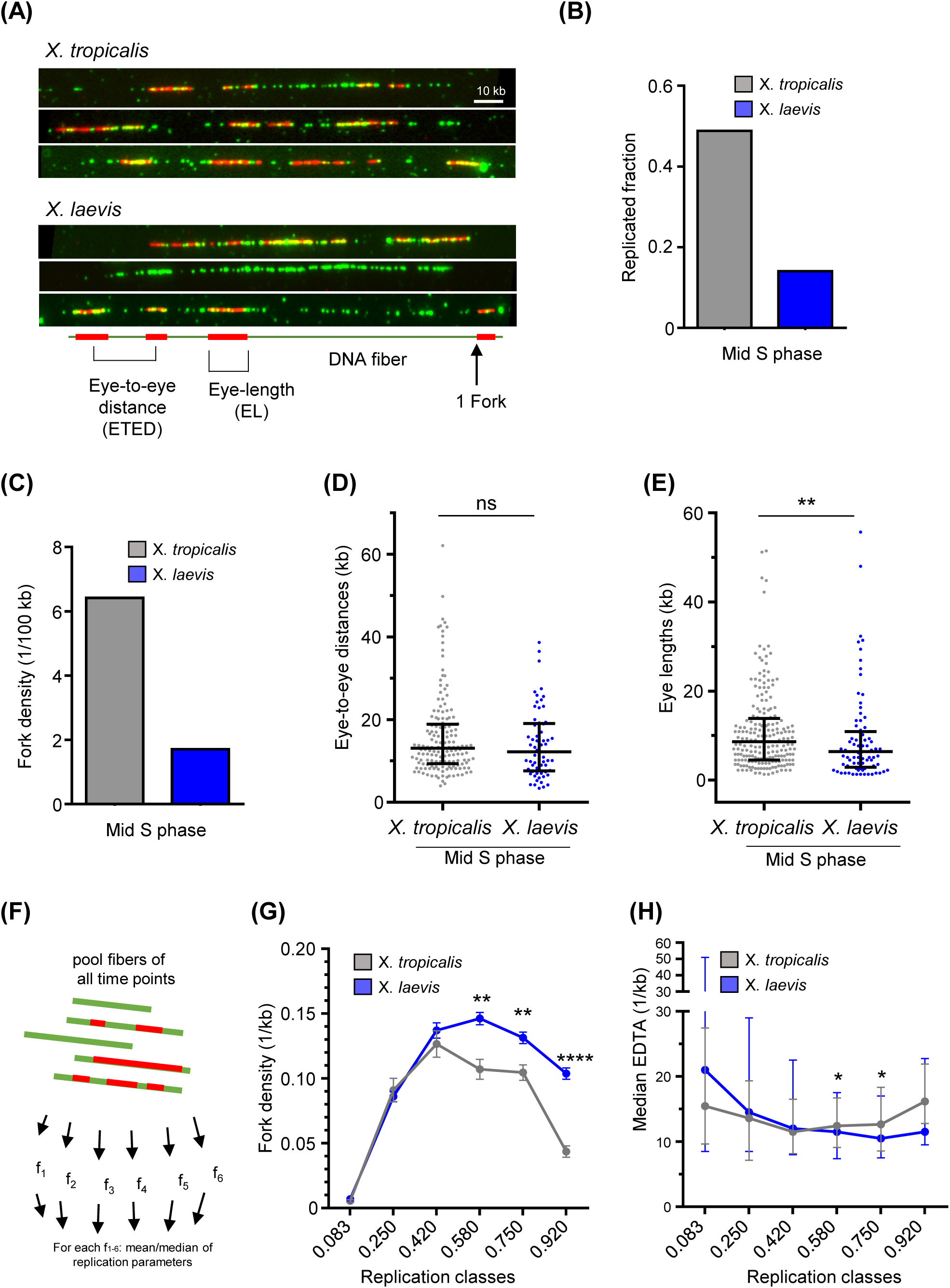
Higher replication fork density and apparent fork speed in *X. tropicalis* than in *X. laevis* sperm nuclei on single DNA molecules at a given time. Sperm nuclei were incubated in the same egg extract as in Fig. 2 with biotin-dUTP, reactions were stopped at 60 and 90 min, and genomic DNA was purified for DNA combing analysis. (A) Representative immunofluorescence-microscopic images of DNA fibers from the 90 min time point = Mid S phase (green: DNA, red: biotin-labeled replication tracks). EL = eye length, ETED = eye-to-eye distance. Scale bar = [10kb]. (B) Replicated fraction at mid S phase. (C) Replication fork density at mid S phase (1 fork/100 kb). (D) Eye-to-eye distances at mid S phase (scatter plot with median and interquartile range). n=160, 65. (E) Eye length at mid S phase (scatter plot with median and interquartile range). n=206, 83. (F) Comparison of *X. tropicalis* and *X. laevis* sperm replication kinetics at identical replication extent by sorting DNA fibers from two independent experiments for *X. tropicalis* (Suppl. Table 1 and 2) and *X. laevis* [35] into six replication extent classes: 0.083[0-0.17], 0.25[0.17-0.33], 0.42[0.33-0.50], 0.58[0.50-0.67], 0.75[0.67-0.83], 0.92[0.83-1]; replication parameter values per class were calculated. (G) Mean fork density with SEM per replication extent class midpoints (H). Median eye-to-eye distances with interquartile range per replication extent class midpoint. * Indicates significant difference (Mann-Whitney U test, two-sided, p<0.05: p-values: * 0.01-0.05; ** 0.001-0.01; *** 0.0001-0.001; ****<0.0001).

### Comparison of replication kinetics in interspecies system between *Xenopus tropicalis* (2x), *Xenopus laevis* (4x), and *Xenopus eysoole* (12x)

We added the dodecaploid *X. eysoole* to our comparative study, and incubated in parallel sperm nuclei from all three species at the same concentration (2000 nuclei/µl) in *X. laevis* egg extract in the presence of rhodamine-dUTP (Fig. 4A). We stopped the reaction after 40 and 60 minutes. Nuclei from all three species decondensed and replicated (Fig. 4B). The nuclear area of replicating *X. eysoole* sperm was significantly larger than that of the two other species (Fig. 4C). We quantified the mean rhodamine fluorescence per nucleus in all three species after size normalization. As shown in Fig. 2 and Fig. 3, we observed a decrease in DNA synthesis associated with increasing polyploidy from 2x to 4x. This decrease was corroborated by a 3.6 and 6-fold reduction in the mean rhodamine fluorescence in *X. laevis* compared to *X. tropicalis* nuclei at 40 and 60 min, respectively (Fig. 4D). We expected that the DNA synthesis would further decrease in dodecaploid *X. eysoole* nuclei compared to *X. laevis*. However, quantification of the rhodamine signal in *X. eysoole* nuclei revealed an 11- and 4-fold higher signal at respectively 40 and 60 min than in *X. laevis* replicating sperm. This increase in DNA synthesis in *X. eysoole* compared to *X. laevis* nuclei was confirmed in a second independent experiment (Fig. 4E). These findings suggest a different coupling of replication dynamics to genome size in the higher dodecaploid sperm species within the interspecies system.

**Fig. 4:**
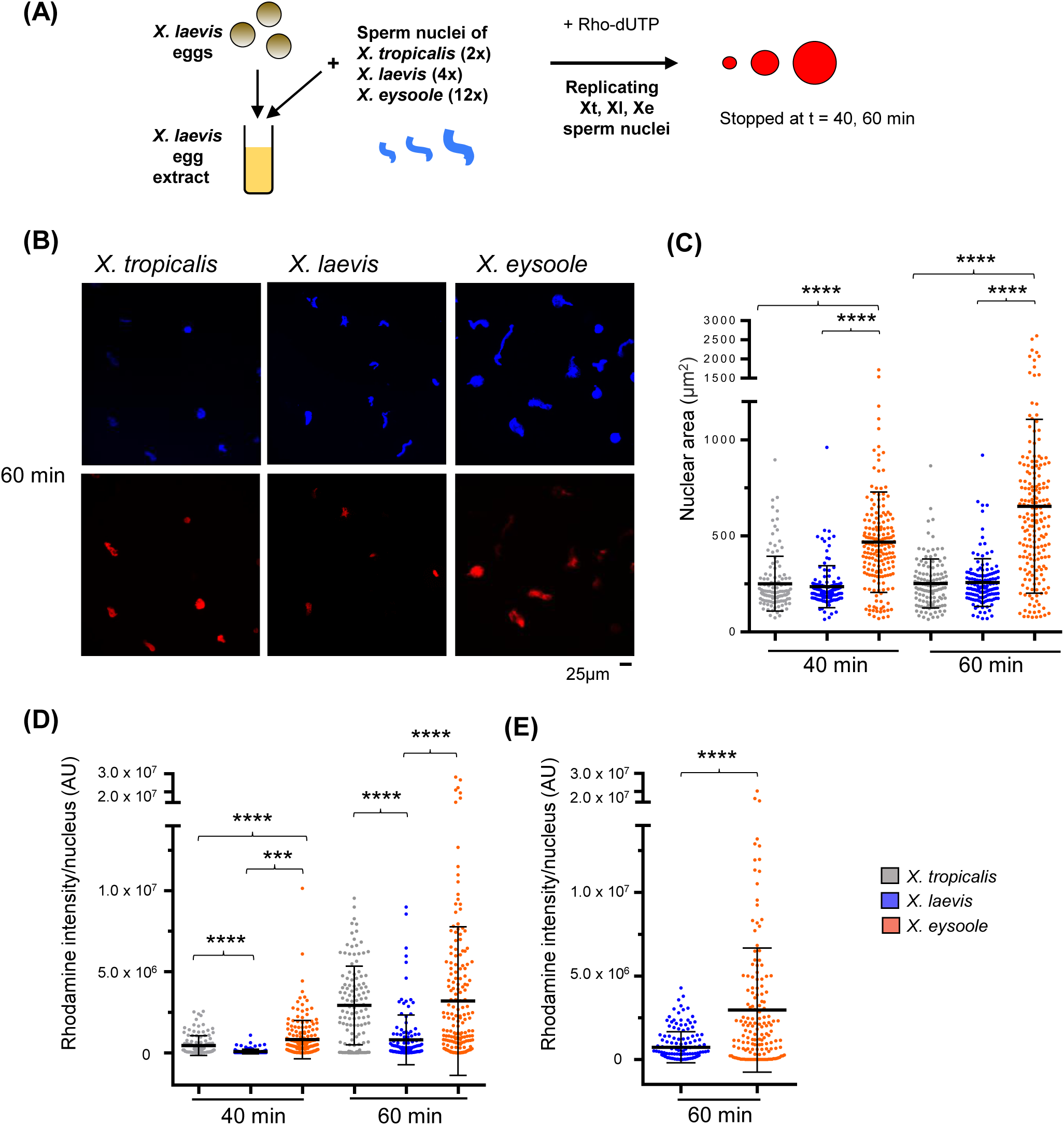
Comparison of nuclear area and rhodamine-dUTP incorporation in three *Xenopus* sperm species replicating in *X. laevis* egg extract. (A) Experimental outline: Interphase egg extract is prepared from *X. laevis* eggs, and sperm nuclei from *X.tropicalis*, *X.laevis,* and *X. eysoole* are incubated in the presence of rhodamine-dUTP for 40min and 60 min. (B) Representative images of sperm nuclei (blue: Hoechst, red: rhodamine) at 60 min. (C) Quantification of nuclear area (µm^2^) (scatter plot with mean ± SD). *n* = [123, 128, 183; 120, 125, 193] at 40min and 60 min. (D) Quantification of rhodamine intensity per nucleus, arbitrary unit (AU) (scatter plot with mean ± SD). *n* = [123, 128, 183; 120, 125, 193]. Scale bar = [25 µm]. (E) Second biological replicate at 60 min, as in (D). Quantification of rhodamine intensity per nucleus, *n* = [154, 151]; * indicates significant difference (Mann-Whitney U test, two-sided, p<0.05: p-values: * 0.01-0.05; ** 0.001-0.01; *** 0.0001-0.001; ****<0.0001).

### Comparative analysis of conservation of replication-related genes in the three *Xenopus* species

In a polyploid, redundant functional elements are expected to rapidly revert to single copies unless prevented by neofunctionalization, subfunctionalization, or selection for gene dosage. We aimed to quantify the extent of gene duplication and protein sequence conservation across *X. tropicalis*, *X. laevis*, and *X. eysoole* to characterize the evolution of replication-related genes following polyploidization events. We established a list of 85 genes directly involved in DNA replication initiation and elongation phases in *Xenopus* and retrieved one reference sequence for each *X. tropicalis* and *X. laevis* ortholog (Suppl. Table 3-4). In addition, *X. eysoole* orthologs were identified from a Trinity assembly generated by RNA-seq (Suppl. Table 5). The allotetraploid *X. laevis* genome is organized into two distinct homoeologous S and L subgenomes, with preferential loss or retention of some functional gene categories, with implications for dosage compensation after WGD [23]. Our analysis revealed that DNA replication-related genes map predominantly to the dominant L chromosomes (54/85, i.e. 64%) and that about two-thirds of replication-related genes underwent rediploidization after allotetraploidization (39/62, i.e. 63%) (Fig. 5A), a higher percentage than what has been observed for the rediploidization of all protein-coding genes during *X. laevis* evolution (43.6%). Replication initiation genes such as Orc1-6, Cdc45, Treslin (Ticrr), Mtbp, MCM10, and DNA polymerases (Pol α, ε, δ) have mainly rediploidized, whereas elongation factors (PCNA, RPA, RFC2-4) and licensing factors involved in helicase loading (Cdc6, Geminin, Cdt1) were retained (Fig. 5B). The replicative helicase MCM2–7 complex presents an intriguing case. Two isoforms of the MCM3 and MCM6 subunits, each showing relatively low conservation (∼71%), are differentially expressed during development, with distinct maternal and zygotic forms [30,31]. Except MCM3—whose two isoforms both reside on the L subgenome—all other MCM homoeologs (MCM2, 4, 5, 6, 7) are retained on both L and S subgenomes and remain highly conserved (>95% identity). Embryonic protein data [32] show that the L-subgenome proteins MCM2, 4, 5, 6, and 7 are expressed from the earliest stages (Fig. 5C). In contrast, expression of their S-subgenome counterparts increases from neurulation onward, paralleling the pattern observed for the two divergent MCM3 isoforms. For MCM6, three proteins are detected: one maternal form from the L subgenome and two zygotic forms from the S and L subgenomes. Thus, the MCM2–7 complex exhibits a coordinated, subgenome-specific, and developmentally regulated expression program.

**Fig. 5:**
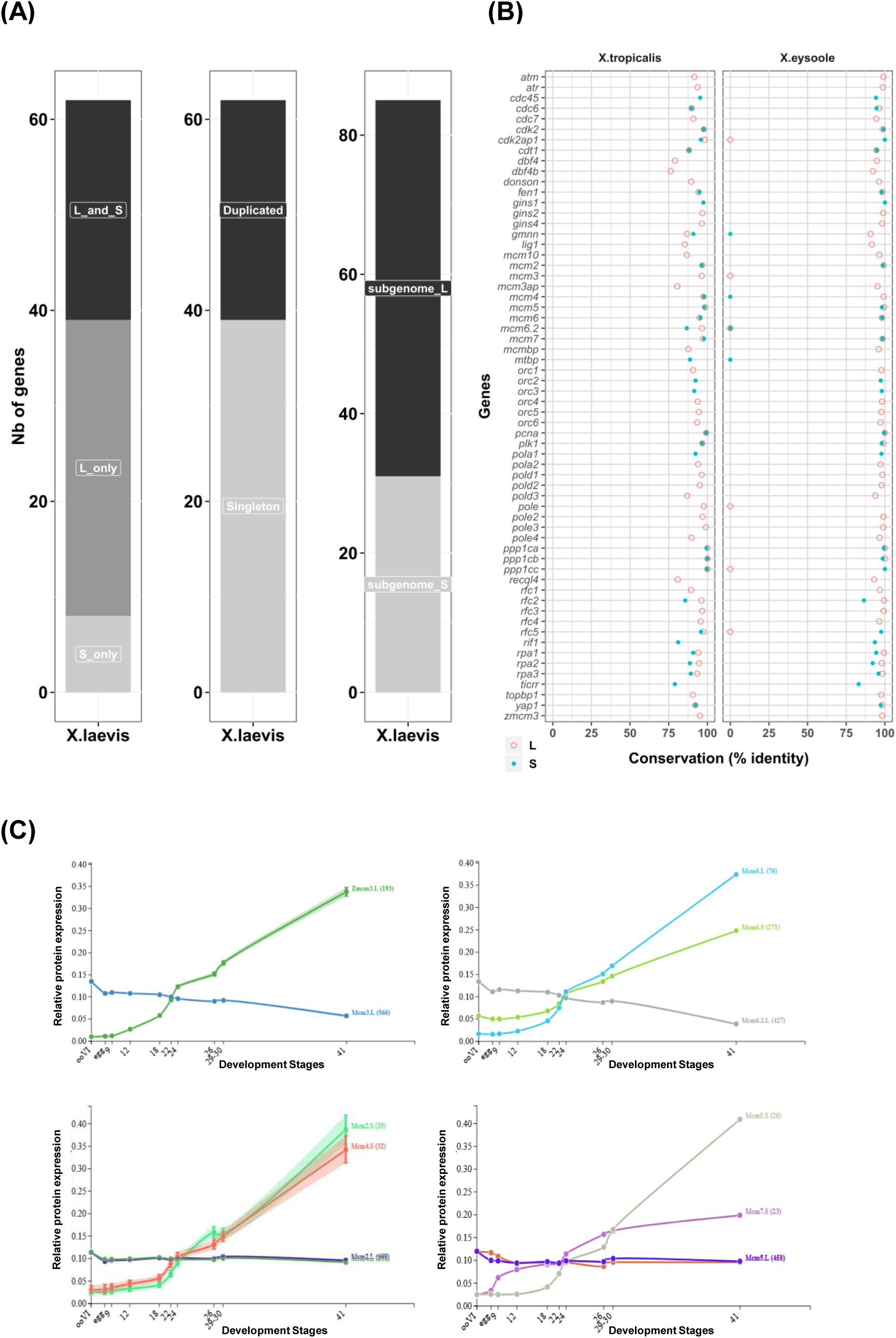
Retention and conservation of replication genes and proteins in *X. tropicalis, X. laevis,* and *X. eysoole*. (A) Subgenome mapping and duplication status of replication genes in the tetraploid *X. laevis* genome, indicating the number of genes in S, L, or S and L subgenomes (left panel), the number of genes retained (duplicated) or diploidized (singletons) (mid panel), the number of genes on L or S genome (right panel). The number of unique (n=62) or total genes (n=85) retained in either L-genome, S-genome, or both subgenomes is indicated. (B) Conservation (% identity) of orthologous *X. laevis* replication proteins in *X. tropicalis* and in *X. eysoole,* shown for L (red circles) and S (blue circles) homoeologous genes in the *X. laevis* genome. (C) Distinct embryonic protein expression of isoforms and homoeologues of the replicative helicase MCM2-7 subunits during early *X. laevis* development (Xenbase (http://www.xenbase.org/, RRID: SCR_003280 using protein data from [32])). zMCM3 and MCM6.L and MCM6.S are distinct zygotic isoforms with lower sequence conservation towards the maternally expressed proteins MCM3L and MCM6.2L. Numbers in brackets indicate the detected peptide number.

In *X. eysoole*, we identified orthologs for 74 out of the 85 *X. laevis* genes (87%), exhibiting the same pattern of duplicated gene retention and no evidence for additional homoeologs (Suppl. Table 5). Protein sequence comparisons showed an extremely high level of conservation between *X. laevis* and *X. eysoole* (median identity: 97.82%), and a very high conservation between *X. laevis* and *X. tropicalis* (median identity: 94.7%; Fig. 5B, Suppl. Table 6-7). Only a few proteins displayed around 80% or less conservation between *X. tropicalis* and *X. laevis*, including the rate-limiting firing factors such as the maternal and zygotic homologs of Dbf4b (Drf1), Dbf4, Treslin (Ticrr), and RecQL4. Altogether, these data indicate that most replication-related proteins are highly conserved across these three *Xenopus* species, suggesting strong evolutionary constraints on the DNA replication machinery despite extensive genome duplication events.

## Discussion

Polyploidy is a major evolutionary force driving genomic restructuring, gene duplication, and speciation across the tree of life [33]. However, how polyploidy impacts DNA replication, a process essential for genome maintenance and stability, remains poorly understood. The *Xenopus* genus, which comprises species with a broad range of ploidy levels, offers a unique system for investigating how genome size shapes the dynamics of DNA replication. Here, we established an interspecies *in vitro* system to mimic a two to six-fold increase in chromosome number while keeping the nuclei concentration and protein factor availability constant, titrating replication factors. We show that replication dynamics scale differently with different genome sizes.

### Different responses of the replication program to an increasing polyploidy

Our single-molecule analysis revealed that a two-fold increase in chromosome number slows DNA replication by globally decreasing origin activation rather than locally increasing inter-origin distances in *X. laevis* nuclei (4x) compared with *X. tropicalis* nuclei (2x). This suggests increased asynchrony in the replication cluster activation, likely extending the temporal program in *X. laevis*, although some compensation at higher replication extents was observed through increased origin activation inside replication clusters. This latter result closely resembles what has been described when the nuclear to cytoplasmic (N/C) concentration ratio is experimentally increased in *X. laevis* egg extracts [13] and, at first sight, supports the model that an elevated DNA content titrates limiting replication factors either *in vivo* at the MBT [14,34] or in extracts [27]. Interestingly, we recently showed that the absence of the replication timing factor Rif1 in *X. laevis* sperm accelerates the replication timing program *in vitro*, likely by facilitating the chromatin access of limiting origin-firing factors, such as Treslin/Mtbp, TopBP1, and Cdc7/Drf1 [35]. The faster replication of *X. tropicalis* sperm nuclei compared to *X. laevis* might therefore mirror the effects of Rif1 depletion. Since the conservation of Rif1 between *X. tropicalis* and *X. laevis* is also amongst the lowest (81%), it is therefore tempting to speculate that Rif1/PP1 activity could be reduced or less efficient on *X. tropicalis*-assembled chromatin. Other critical factors are also less conserved, such as the firing factors Dbf4, a subunit of the S-phase kinase DDK, Treslin, and RecQL4. Future structure-function experiments will be required to test whether specific changes in protein sequences resulted in adaptation of the replication program to different polyploid statuses.

When we compared replication kinetics across the three species with different ploidy levels, we found that *X.eysoole* (12x) nuclei replicated markedly faster than *X. laevis* nuclei. This observation argues against the hypothesis that titrating the replication-limiting factors would limit replication kinetics in the higher polyploid species in the interspecies *in vitro* system. An alternative explanation is that different sperm DNA templates exhibit species-specific replication efficiencies. Although *Xenopus laevis* egg extracts are known to efficiently replicate any added DNA template, *X. eysoole* sperm, and to a lesser extent *X. tropicalis* sperm, may assemble into a more replication-permissive chromatin, promoting rapid recruitment of replication factors and origin firing by S-phase kinases. However, so far, no DNA sequence-specific elements are known to control origin activation during early *Xenopus* embryogenesis [7]. We propose that chromatin context or dynamics, nuclear scaling, or the availability of regulatory factors may play a greater role than sequence variation in these species-specific differences.

A recent study using *X. longipes*, another dodecaploid species, showed that early embryogenesis proceeds almost identically to that of *X. laevis* until MBT and then slows down at neurulation (stage 13, ∼16 hpf) [33]. MBT occurred at a lower N/C ratio *in X. laevis* than in *X. tropicalis* or *X. longipes*, suggesting that developmental timing scales non-linearly with genome size and possibly with replication capacity, as observed in our study. However, DNA replication was not investigated in this study. Interspecies *Xenopu*s extract systems have been successfully used to study nuclear scaling [41] and spindle assembly [42,43], in mitotic or cycling extracts. Brown *et al*. showed that titratable nuclear import factors have a greater influence on nuclear size than DNA content. Additionally, mitotic spindles are smaller when assembled in *X. tropicalis* extract compared to *X. laevis* extract, and factors involved in spindle assembly regulation vary between *X. laevis* and *X. tropicalis* extracts. Exploring whether nuclear import regulation could explain our observations would be intriguing; however, we did not observe differences in sperm decondensation or in the start of S phase between the nuclei of different species. Since these steps depend on the import of chromatin and replication factors in this *in vitro* system, this suggests that nuclear import does not play a major role in our results, at least regarding the initiation of S phase.

### Genome response to increasing polyploidy for DNA replication genes

From an evolutionary standpoint, the coordinated expression of gene products within macromolecular complexes in defined stoichiometric amounts strongly influences the evolutionary trajectory of duplicated genes [36]. In this study, we found that only one-third of replication gene homoeologs remain duplicated, including the replicative polymerases and often those found in protein complexes (MCM2-7, RPA1-3). An interesting example is the replicative helicase MCM2-7 complex, which plays a central role in the regulation of origin firing. We observed that all subunits, except MCM3, are retained but differentially expressed, suggesting a sub-functionalization of MCM subunits during development, mainly on the S-genome, after genome duplication. The differential expression of a distinct zygotic MCM2-7 complex could contribute to the change in origin usage and the slower replication program observed after MBT [14,15], but detailed structure-function analysis is required to understand this phenomenon better. The low overall retention rate indicates that dosage compensation mechanisms have operated on these replication factors following genome duplication in *X. laevis,* thereby regulating their abundance to preserve genome stability and proper cell cycle progression.

On the other hand, we observed that among the factors whose genes underwent rediploidization are all initiation factors, such as Orc1-6, and, surprisingly, also the low-abundant, rate-limiting factors for origin activation, such as Treslin (Ticcr), RecQL4, Cdc45, TopBP1, and Dbf4. This suggests that, for these initiation factors, either no dosage selection occurred, single loci encoding proteins are sufficient, or homoeologous genes may be functionally incompatible. These observations warrant future quantitative and comparative analysis of replication factor concentrations in *X. tropicalis*, *X. laevis*, and *X. eysoole* eggs and embryos, which may elucidate how S-phase regulation adapts to genome size. Future genomic and transcriptomic studies of *X. eysoole* will further illuminate how replication-related genes, replication programs, and cell proliferation evolved in higher polyploid *Xenopus* species. Beyond *Xenopus*, polyploidization events across plants and yeasts frequently involve rebalancing of transcriptional networks to maintain cell viability [37]. The ability of highly polyploid species to sustain efficient S-phase progression without proportional increases in replication factor abundance suggests the evolution of compensatory regulatory mechanisms, such as altered replication timing control, altered chromatin accessibility, or enhanced replication origin density.

It is important to acknowledge the limitations of our *in vitro* approach. While *Xenopus* egg extracts provide a robust system for dissecting replication mechanisms under controlled conditions, they may not fully recapitulate the native chromatin organization, developmental regulation, or spatial constraints found in the embryo. Future *in vivo* analyses of replication dynamics and factor localization in polyploid *Xenopus* species will be crucial for validating whether the mechanisms inferred from extract systems operate during early embryonic development.

In conclusion, our findings suggest that replication kinetics can, but do not necessarily, slow down with increasing genome size and that species-specific adaptations, together with replication-factor dosage, buffer replication efficiency as high ploidy is reached. This work establishes *Xenopus* as a tractable model for investigating how polyploidy reshapes replication programs and provides an evolutionary framework for understanding genome stability in species with expanded genomic content. It may also offer insights into how the unscheduled whole-genome duplications observed in many human tumors enhance their adaptive potential, particularly during later stages of cancer evolution [38,39].

## Materials and Methods

### Ethics statement

All animal experiments have been carried out in accordance with the European Community Council Directive of 22 September 2010 (2010/63/EEC). All animal care and experimentation were conducted according to institutional guidelines under the institutional license C 91-471-102 at the CNRS aquatic animal facility in Gif-sur-Yvette. The institutional animal care committee CEEA #59 approved the study protocols, and we received an authorization from the Direction Départementale de la Protection des Populations under the reference APAFIS#998-2015062510022908v2 for *Xenopus* experiments. *X. eysoole* were collected during field research in Oku conducted under the permissions issued by the local traditional authorities and by the Cameroon Ministry of Scientific Research and Innovation (MINRESI:34/MINRESI/B00/C00/C10/C12) and the Ministry of Forestry and Wildlife (MINFOF:1235/PRS/MINFOF/SETAT/SG/DFAP/SDVEF/SC/BJ) and exported under the CITES permit 0384/P/MINFOF/SG/DFAP/DVEF/SC. All *X. eysoole* frogs were bred or humanely euthanized following the procedures defined by the EGCE aquatic animal facility of University Paris-Saclay (French government authorization APAFIS#43700-2023060111493334-v4 following the ethical committee of Paris Centre et Sud; authorization by the Préfecture de l’Essonne A91-272-111).

### Replication of sperm nuclei in *Xenopus* egg extracts

Replication-competent extracts from unfertilized *Xenopus* eggs and sperm nuclei from the testes of male frogs from all species were prepared as described [6]. Sperm nuclei from *X. laevis, X. tropicalis, and X. eysoole* (2000 nuclei/µl) were incubated in *X. laevis* interphase egg extract in the presence of cycloheximide to inhibit translation (250 µg/ml, Sigma), energy mix (7.5 mM creatine phosphate, 1 mM ATP, 0.1 mM EGTA, pH 7.7, 1 mM MgCl_2_). Reactions were stopped at indicated times and treated for fluorescence microscopy, DNA gel electrophoresis, or DNA combing.

### Neutral and alkaline agarose gel electrophoresis

Sperm nuclei were incubated in fresh extracts complemented with indicated reagents and one-fiftieth volume of α-[^32^P]-dCTP (3000 Ci/mmol). DNA was recovered after DNAzol® treatment (Invitrogen protocol), ethanol precipitation, separated on 1.1% alkaline agarose gels, and analyzed as described [10].

### Molecular combing of DNA and detection by fluorescent antibodies

Biotin-labelled sperm DNA was extracted and combed as described [40]. Biotin was detected with AlexaFluor594-conjugated streptavidin, followed by anti-avidin biotinylated antibodies. This was repeated twice, followed by mouse ssDNA antibody, Alexa Fluor 488 rabbit anti-mouse, and Alexa Fluor 488 goat anti-rabbit for enhancement [41]. Images of the combed DNA molecules were acquired and measured as described [40]. The fields of view were chosen at random. Measurements on each molecule were made using Fiji software [42] and compiled using macros in Microsoft Excel. Replication eyes were defined as the incorporation tracks of biotin–dUTP. Replication eyes were considered to be the result of two replication forks, and incorporation tracks at the extremities of DNA fibers were considered to be the products of one replication fork. Tracts of biotin-labelled DNA needed to be at least 1 kb in length to be considered significant and scored as eyes. When the label was discontinuous, the unlabeled DNA tract was considered a real gap if it was more than 1 kb in length. The replicated fraction of each fiber was calculated as the sum of eye lengths (red tracks) divided by the total DNA length (green track). Fork density was calculated as the total DNA divided by the total number of forks. The midpoints of replication eyes were defined as the origins of replication. Eye-to-eye distances (ETED), also known as inter-origin distances, were measured between the midpoints of adjacent replication eyes. Incorporation tracks at the extremities of DNA fibers were not regarded as replication eyes. Still, they were included in determining the extent of replication, or the replicated fraction, which was calculated as the sum of all eye lengths (EL) divided by the total DNA.

### Fluorescent analysis of replicating nuclei

Sperm nuclei (2000/μl) were added to replication reactions in the presence of 20 μM rhodamine-dUTP (Roche) and stopped at indicated time points or pulse labeled for the stated time period; 20 μl aliquots were diluted in 500 μl PBS and fixed by the addition of 500 μl p-formaldehyde 8%. Nuclei were spun onto coverslips using a 1 ml 20% sucrose cushion in PBS. After washing, coverslips were incubated with Hoechst 33258. After washing, mounted coverslips were imaged for Hoechst and rhodamine, using an Olympus BX63 fluorescence microscope, and area size and rhodamine fluorescence intensity/nucleus were quantified as described [43]. Using Analyze Particles of the Fiji software [42], areas of Hoechst-stained nuclei were saved as ROI (region of interest). The rhodamine staining intensity in each ROI and the background of each slide were measured. For each nucleus, Corrected Total Fluorescence (CTF) was calculated using the following equations: CTF = Integrated Density of selected nuclei - (Area of selected nuclei X Mean fluorescence of background).

### Transcriptomics in *X. eysoole*

We obtained embryos from natural matings of *X. eysoole* and prepared total RNA from tailbud-stage embryos (NF30) as well as from tadpole tails and heads (NF55) using standard protocols. We then purified mRNA using the NEBNext polyA mRNA magnetic module kit (New England Biolabs) [44]. The I2BC sequencing facility prepared RNAseq libraries from these polyA+ RNA using an mRNA-stranded approach, followed by sequencing using a NextSeq 500/550 Mid output kit v2 (75 cycles). We obtained a total of 58.5, 60.8, and 56.0 million reads after post-adapter trimming and filtering for the NF55 tail, NF30, and NF55 head samples, respectively. We filtered out reads matching *X. laevis* mitochondrial and ribosomal RNA transcripts, aligned them to the *X. laevis* genome assembly (v10_1; GCA_017654675.1) using HiSat2, and then ran Trinity with a genome-guided assembly.

### Analysis of gene duplication and replication protein sequence conservation across *X. tropicalis*, *X. laevis*, and *X. eysoole*

We listed 85 *Xenopus* genes that play crucial roles in replication initiation or elongation and extracted their corresponding nucleotide and protein accession numbers for *X. laevis* and *X. tropicalis* using Xenbase [45] and the NCBI Datasets tools. We selected either the RefSeq entry or the longest isoform of the *X. laevis* protein sequences and compared them to their *X. tropicalis* ortholog or to the *X. eysoole* Trinity assembly using tblastn. We manually curated the alignments to assess L and S gene orthology between *X. laevis* and *X. eysoole* and between *X. laevis* and *X. tropicalis* to build the data tables provided in Suppl. data. MCM2-7 embryonic expression graphs were generated by Xenbase [46] (http://www.xenbase.org/, RRID:SCR_003280) using protein data from [32].

### Statistics and reproducibility

Statistical analysis was performed using GraphPad Prism software, version 8.3.0, with p-values and sample sizes reported in the figure legends. A p-value **≤**0.05 was considered significant.

## Data availability

The datasets analyzed in the current study are available from the corresponding author upon request. The *X. eysoole* RNA-seq data and transcriptome assembly for this study have been deposited in the European Nucleotide Archive (ENA) at EMBL-EBI under accession number PRJEB104886 (https://www.ebi.ac.uk/ena/browser/view/PRJEB104886).

## Supporting information

Suupl. Data

## Supplementary data

Supplementary Tables 1-2: DNA combing data summaries for Fig. 3.

Supplementary tables 3-7: Tables of replication gene comparisons, subgenome expression, and gene sequence conservation for Fig. 5.

## Acknowledgements

We thank Leonid Peshkin for the discussions and Marie-Noëlle Prioleau for the critical reading of the manuscript. This work was supported by a China Scholarship Council – Université Paris Saclay PhD fellowship to Z.L. (202106760020).

## Author contributions

H.N., O.H., Z.L., N.P., and K.M. performed and analyzed experiments. K.M. wrote the manuscript with the help of N.P. and O.H.

## Abbreviations

Cdc6: Cell division cycle 6
Cdc7: Cell division cycle 7
Cdc45: Cell division cycle 45
Cdk: Cyclin-dependent kinase
Cdt1: Cdc10-dependent transcript 1
CMG: Cdc45-MCM-GINS
CTF: Corrected total fluorescence
Dbf4: Dumbbell former 4 protein
dCTP: deoxycytidine triphosphate
DDK: Dbf4-dependent kinase
dUTP: deoxyuridine triphosphate
DNA: Deoxyribonucleic acid
Drf1: DBF4-related factor 1
Donson: Downstream Neighbor of Son
GINS: Go-ichi-ni-san (5-1-2-3 japanese, Sld5, PSf1, Psf2, PSf3)
G1 phase: Gap phase 1
kbp: kilobase pairs
kDa: kilo Dalton
MBT: Mid-blastula transition
MCM: Minichromosome maintenance
MTBP: Mdm2-binding protein
N/C ratio: Nucleo-cytosolic ratio
ORC: Origin recognition complex
Pre-RC: Pre-replication complex
Pre-IC: Pre-initiation complex
PP1: Protein phosphatase 1
PCNA: Proliferating cell nuclear antigen
RecQL4: RecQ-like helicase 4
RFC: Replication factor C
Rif1: Rap1 interaction factor
RPA: Replication protein A
RNA: Ribonucleic acid
ROI: Region of interest
S phase: Synthesis phase
TICCR: TopBP1 interacting checkpoint and replication regulator
TopBP1: DNA topoisomerase binding protein 1
WGD: Whole genome duplication
X. laevis: Xenopus laevis
X. tropicalis: Xenopus tropicalis
X. eysoole: Xenopus eysoole

