## Supplementary material for "Ploidy-dependent modulation of DNA replication kinetics in *Xenopus*": Suupl. Data

Supplementary Tables 1-2: DNA combing data summaries for Fig. 3.

Supplementary Tables 3-7: Tables of replication gene comparisons, subgenome expression, and gene sequence conservation for Fig. 5.

**Supplementary Table 1 : DNA combing data summary from Fig. 3 A-F.** *X. laevis* and *X. tropicalis* sperm replicates at 60 and 90 min in interphase *X. laevis* egg extracts.

| **sperm nuclei (2000 n/µl)** | ***X. tropicalis*** | ***X. tropicalis*** | ***X. laevis*** | ***X. laevis*** |
|---|---|---|---|---|
| | 60 min | 90 min | 60 min | 90 min |
| Analysed DNA (kb) | 8012.80 | 7267.20 | 9562.56 | 11238.08 |
| Replicated DNA(kb) | 2576.64 | 3547.84 | 525.76 | 1579.20 |
| Replication Extent | 0.32 | 0.49 | 0.05 | 0.14 |
| Number of analysed fibers | 88 | 75 | 91 | 112 |
| Number of fully replicated fibers | 2 | 2 | 0 | 6 |
| Number of unreplicated Fibers | 48 | 21 | 73 | 83 |
| Average size of all fibers (kb) | 91.05 | 96.90 | 105.08 | 100.34 |
| Average size of fully replicated fibers (kb) | 110.40 | 31.52 | 0.00 | 56.64 |
| Average size of unreplicated fibers (kb) | 82.05 | 91.40 | 102.70 | 102.62 |
| Number of replication eyes | 120 | 206 | 75 | 83 |
| Average Eye Length (kb) | | | | |
| Local | 10.34 | 11.73 | 6.00 | 9.27 |
| Overall | 18.40 | 15.19 | 6.74 | 16.45 |
| Number of eye to eye distances (ETED) | 89 | 160 | 57 | 65 |
| Average ETED length (kb) | | | | |
| Local | 17.02 | 15.99 | 14.48 | 15.15 |
| Overall | 57.23 | 31.12 | 122.60 | 117.06 |
| Number of replication fork | 280 | 467 | 156 | 192 |
| Fork Density (forks/100 kb) | 3.49 | 6.43 | 1.63 | 1.71 |

**Supplementary Table 2 : DNA combing data replicate *for X. tropicalis* for Fig. 3 G.H .** *X. tropicalis* sperm replicates at indicated times in interphase *X. laevis* egg extracts.

| ***X. tropicalis*** | 35min | 40min | 50min |
|---|---|---|---|
| Analysed DNA (kb) | 5822.0 | 4902.4 | 6676.0 |
| Replicated DNA(kb) | 597.5 | 986.6 | 1801.5 |
| Replication Extent | 0.103 | 0.201 | 0.270 |
| Number of analysed fibers | 104 | 89 | 126 |
| Number of fully replicated fibers | 0 | 1 | 1 |
| Number of unreplicated Fibers | 74 | 53 | 56 |
| Average size of all fibers (kb) | 56.0 | 55.1 | 53.0 |
| Average size of fully replicated fibers (kb) | 0.0 | 52.4 | 37.4 |
| Average size of unreplicated fibers (kb) | 56.2 | 54.2 | 51.1 |
| Number of replication eyes | 68 | 57 | 186 |
| Average Eye Length (kb) : | | | |
| Local | 6.3 | 8.9 | 6.0 |
| Overall | 7.5 | 13.0 | 8.3 |
| Number of eye to eye distances (ETED) | 41 | 31 | 123 |
| Average ETED length (kb) : | | | |
| Local | 11.4 | 13.2 | 12.0 |
| Overall | 73.2 | 64.5 | 30.8 |
| Number of replication fork | 159 | 152 | 433 |
| Fork Density (forks/100 kb) | 2.73 | 3.10 | 6.49 |

### Supplementary Table 3: Replication genes *X. tropicalis*

| NCBI GeneID | Symbol | Transcript Accession | Transcript Length | Transcript Protein Accession | Transcript Protein Length | Transcript Protein Name | Transcript Protein Isoform |
|---|---|---|---|---|---|---|---|
| 100490767 | atm | XM_002934928.5 | 10039 | XP_002934974.3 | 3060 | serine-protein kinase ATM | isoform X1 |
| 100495593 | atr | XM_031902384.1 | 8400 | XP_031758244.1 | 2655 | serine/threonine-protein kinase ATR | isoform X1 |
| 779545 | cdc45 | XM_002931882.5 | 2027 | XP_002931928.1 | 569 | cell division control protein 45 homolog | |
| 493356 | cdc6 | NM_001007993.1 | 2232 | NP_001007994.1 | 554 | cell division control protein 6 homolog | |
| 549335 | cdc7 | NM_001016581.2 | 1687 | NP_001016581.1 | 483 | cell division cycle 7-related protein kinase | |
| 493498 | cdk2 | NM_001008135.1 | 1903 | NP_001008136.1 | 297 | cyclin-dependent kinase 2 | |
| 549484 | cdk2ap1 | NM_001016730.2 | 1238 | NP_001016730.1 | 116 | cyclin-dependent kinase 2-associated protein 1 | |
| 100379812 | cdt1 | XM_002933654.5 | 2717 | XP_002933700.2 | 617 | DNA replication factor Cdt1 | |
| 733748 | dbf4 | NM_001044505.1 | 2493 | NP_001037970.1 | 663 | protein DBF4 homolog A | |
| 100379975 | dbf4b | XM_004918590.4 | 4123 | XP_004918647.2 | 809 | protein DBF4 homolog B | isoform X1 |
| 395018 | donson | NM_204053.1 | 2399 | NP_989384.1 | 577 | protein downstream neighbor of son homolog | |
| 549759 | fen1 | NM_001017005.2 | 1436 | NP_001017005.1 | 382 | flap endonuclease 1 | |
| 548918 | gins1 | NM_001016164.2 | 752 | NP_001016164.1 | 196 | DNA replication complex GINS protein PSF1 | |
| 549769 | gins2 | NM_001017015.2 | 1483 | NP_001017015.1 | 185 | DNA replication complex GINS protein PSF2 | |
| 496455 | gins4 | NM_001011045.1 | 1744 | NP_001011045.1 | 221 | DNA replication complex GINS protein SLD5 | |
| 496859 | gmnn | NM_001039736.1 | 1047 | NP_001034825.1 | 219 | geminin | |
| 100271763 | lig1 | XM_002941883.5 | 4372 | XP_002941929.1 | 1040 | DNA ligase 1 | |
| 549663 | mcm10 | NM_001016909.2 | 3072 | NP_001016909.1 | 845 | protein MCM10 homolog | |
| 448458 | mcm2 | NM_001006771.1 | 3268 | NP_001006772.1 | 884 | DNA replication licensing factor mcm2 | |
| 734092 | mcm3 | XM_002934119.5 | 2578 | XP_002934165.2 | 807 | maternal DNA replication licensing factor mcm3 isoform X2 [Xenopus tropicalis] | |
| 100490007 | mcm3ap | XM_004917891.2 | 8220 | XP_004917948.2 | 2384 | germinal-center associated nuclear protein | isoform X1 |
| 448137 | mcm4 | NM_001005655.1 | 3290 | NP_001005655.1 | 863 | DNA replication licensing factor mcm4 | |
| 550081 | mcm5 | NM_001017327.3 | 3028 | NP_001017327.2 | 735 | DNA replication licensing factor mcm5 | |
| 395030 | mcm6 | NM_204062.1 | 2893 | NP_989393.1 | 823 | zygotic DNA replication licensing factor mcm6 | |
| 548975 | mcm6.2 | NM_001016221.2 | 2625 | NP_001016221.1 | 821 | maternal DNA replication licensing factor mcm6 | |
| 407945 | mcm7 | NM_213712.1 | 2499 | NP_998877.1 | 720 | DNA replication licensing factor mcm7 | |
| 493445 | mcmbp | NM_001008082.1 | 2250 | NP_001008083.1 | 627 | mini-chromosome maintenance complex-binding protein | |
| 779505 | mtbp | NM_001114218.1 | 3195 | NP_001107690.1 | 867 | mdm2-binding protein | |
| 734048 | orc1 | NM_001045729.1 | 2859 | NP_001039194.1 | 888 | origin recognition complex subunit 1 | |
| 780267 | orc2 | NM_001079338.1 | 2016 | NP_001072806.1 | 558 | origin recognition complex subunit 2 | |
| 100125181 | orc3 | NM_001103067.1 | 2257 | NP_001096537.1 | 709 | origin recognition complex subunit 3 | |
| 448135 | orc4 | NM_001005654.1 | 1557 | NP_001005654.1 | 432 | origin recognition complex subunit 4 | |
| 549767 | orc5 | NM_001017013.2 | 1815 | NP_001017013.1 | 448 | origin recognition complex subunit 5 | |
| 549939 | orc6 | NM_001017185.2 | 1148 | NP_001017185.1 | 252 | origin recognition complex subunit 6 | |
| 493302 | pcna | NM_001007920.1 | 1134 | NP_001007921.1 | 261 | proliferating cell nuclear antigen | |
| 407937 | plk1 | NM_213679.2 | 2178 | NP_998844.1 | 598 | serine/threonine-protein kinase PLK1 | |
| 100379926 | pola1 | XM_031896702.1 | 4536 | XP_031752562.1 | 1489 | DNA polymerase alpha catalytic subunit | isoform X1 |
| 448613 | pola2 | NM_001005060.1 | 2765 | NP_001005060.1 | 597 | DNA polymerase alpha subunit B | |
| 734063 | pold1 | NM_001045739.1 | 3993 | NP_001039204.1 | 1109 | DNA polymerase delta catalytic subunit | |
| 448271 | pold2 | NM_001005794.1 | 1701 | NP_001005794.1 | 463 | DNA polymerase delta subunit 2 | |
| 549310 | pold3 | NM_001016556.2 | 1314 | NP_001016556.1 | 214 | DNA polymerase delta subunit 3 | |
| 100498309 | pole | XM_004910635.4 | 7100 | XP_004910692.2 | 2286 | DNA polymerase epsilon catalytic subunit A | |
| 595029 | pole2 | NM_001030470.1 | 1718 | NP_001025641.1 | 527 | DNA polymerase epsilon subunit 2 | |
| 550018 | pole3 | NM_001017264.2 | 829 | NP_001017264.1 | 147 | DNA polymerase epsilon subunit 3 | |
| 549359 | pole4 | NM_001016605.2 | 1085 | NP_001016605.1 | 115 | DNA polymerase epsilon subunit 4 | |
| 407879 | ppp1cc | NM_213670.2 | 1993 | NP_998835.1 | 323 | serine/threonine-protein phosphatase PP1-gamma catalytic subunit | |
| 496958 | ppp1cb | NM_001011467.1 | 1183 | NP_001011467.1 | 327 | serine/threonine-protein phosphatase PP1-beta catalytic subunit | |
| 100145598 | ppp1ca | NM_001127010.1 | 2141 | NP_001120482.1 | 326 | protein phosphatase 1, catalytic subunit, alpha isozyme | |
| 780344 | recql4 | NM_001079414.1 | 5033 | NP_001072882.1 | 1520 | ATP-dependent DNA helicase Q4 | |
| 733476 | rfc1 | NM_001199567.1 | 3988 | NP_001186496.1 | 1144 | replication factor C subunit 1 | |
| 100216026 | rfc2 | NM_001142016.1 | 1182 | NP_001135488.1 | 345 | replication factor C subunit 2 | |
| 100493161 | rfc3 | XM_002934033.5 | 1802 | XP_002934079.1 | 356 | replication factor C subunit 3 | |
| 549117 | rfc4 | NM_001016363.2 | 1214 | NP_001016363.1 | 360 | replication factor C subunit 4 | |
| 496525 | rfc5 | NM_001011112.2 | 1895 | NP_001011112.1 | 335 | replication factor C subunit 5 | |
| 100491835 | rif1 | XM_004917657.4 | 8503 | XP_004917714.2 | 2359 | telomere-associated protein RIF1 | isoform X1 |
| 548449 | rpa1 | NM_001015732.1 | 2979 | NP_001015732.1 | 609 | replication protein A 70 kDa DNA-binding subunit | |
| 448500 | rpa2 | NM_001006794.1 | 1221 | NP_001006795.1 | 275 | replication protein A 32 kDa subunit | |
| 549380 | rpa3 | NM_001016626.2 | 856 | NP_001016626.1 | 121 | replication protein A 14 kDa subunit | |
| 100489448 | ticrr | XM_031899228.1 | 8000 | XP_031755088.1 | 1967 | treslin | |
| 100380063 | topbp1 | XM_002937745.4 | 5547 | XP_002937791.3 | 1515 | DNA topoisomerase 2-binding protein 1 | isoform X1 |
| 100488779 | yap1 | NM_001195768.1 | 1371 | NP_001182697.1 | 456 | transcriptional coactivator YAP1 | |
| 100158601 | zmcm3 | NM_001045766.1 | 3121 | NP_001039231.1 | 809 | zygotic DNA replication licensing factor mcm3 | isoform X1 |

### Supplementary Table 4: Replication genes *X. laevis*

| NCBI GeneID | Symbol | Transcript Accession | Transcript Length | Transcript Protein Accession | Transcript Protein Length | Transcript Protein Name | Transcript Protein Isoform |
|---|---|---|---|---|---|---|---|
| 398148 | atm.L | NM_001088499.1 | 9186 | NP_001081968.1 | 3061 | ATM serine/threonine kinase L homeolog | |
| 398197 | atr.L | NM_001088580.1 | 8346 | NP_001082049.1 | 2655 | serine/threonine-protein kinase atr | |
| 446991 | cdc45.S | NM_001093633.1 | 1967 | NP_001087102.1 | 567 | cell division cycle 45 S homeolog | |
| 403388 | cdc6.L | NM_001090971.1 | 2213 | NP_001084440.1 | 554 | cell division cycle 6 L homeolog | |
| 398082 | cdc6.S | NM_001088375.1 | 2323 | NP_001081844.1 | 554 | cell division cycle 6 S homeolog | |
| 398099 | cdc7.L | NM_001088409.1 | 1665 | NP_001081878.1 | 483 | cell division cycle 7 L homeolog | |
| 432036 | cdk2.L | NM_001091507.1 | 2391 | NP_001084976.1 | 297 | Cyclin-dependent kinase 2-like | |
| 399314 | cdk2.S | NM_001090651.1 | 919 | NP_001084120.1 | 297 | cyclin-dependent kinase 2 | |
| 108716575 | cdk2ap1.L | XM_018262805.2 | 1695 | XP_018118294.1 | 120 | cyclin-dependent kinase 2-associated protein 1 | isoform X1 |
| 380551 | cdk2ap1.S | XM_041580351.1 | 1640 | XP_041436285.1 | 120 | cyclin-dependent kinase 2-associated protein 1 | isoform X1 |
| 398024 | cdt1.L | NM_001088269.1 | 2912 | NP_001081738.1 | 620 | DNA replication factor Cdt1 | |
| 108715404 | cdt1.S | XM_018260503.2 | 3681 | XP_018115992.1 | 618 | DNA replication factor Cdt1-like | isoform X1 |
| 398599 | dbf4.L | NM_001089137.1 | 2336 | NP_001082606.1 | 661 | protein DBF4 homolog A | |
| 398696 | dbf4b.L | NM_001089271.1 | 3811 | NP_001082740.1 | 784 | protein DBF4 homolog B | |
| 495432 | donson.L | NM_001095087.1 | 2768 | NP_001088556.1 | 579 | protein downstream neighbor of son homolog | |
| 394303 | fen1.L | NM_001087491.1 | 1357 | NP_001080960.1 | 382 | flap endonuclease 1-A | |
| 394311 | fen1.S | NM_001087515.1 | 1469 | NP_001080984.1 | 382 | flap endonuclease 1-B | |
| 398096 | gmnn.L | NM_001088403.1 | 1047 | NP_001081872.1 | 216 | geminin, DNA replication inhibitor L homeolog | |
| 399373 | gmnn.S | NM_001090747.1 | 1096 | NP_001084216.1 | 219 | geminin, DNA replication inhibitor S homeolog | |
| 443928 | gins1.S | NM_001092033.1 | 1056 | NP_001085502.1 | 196 | DNA replication complex GINS protein PSF1 | |
| 444053 | gins2.L | NM_001092158.1 | 1084 | NP_001085627.1 | 185 | GINS complex subunit 2 (Psf2 homolog) L homeolog | |
| 414663 | gins4.L | NM_001091233.1 | 1072 | NP_001084702.1 | 221 | DNA replication complex GINS protein SLD5 | |
| 397978 | lig1.L | NM_001088184.1 | 3770 | NP_001081653.1 | 1070 | DNA ligase 1 | |
| 398196 | mcm10.L | NM_001088578.1 | 3497 | NP_001082047.1 | 860 | protein MCM10 homolog | |
| 380451 | mcm2.L | NM_001087290.2 | 3278 | NP_001080759.1 | 886 | DNA replication licensing factor mcm2 | |
| 108715875 | mcm2.S | XM_018261393.2 | 3644 | XP_018116882.1 | 885 | DNA replication licensing factor mcm2-like | isoform X1 |
| 397821 | mcm3.L | NM_001087943.1 | 2548 | NP_001081412.1 | 807 | maternal DNA replication licensing factor mcm3 | |
| 108701406 | mcm3ap.L | XM_041576444.1 | 8512 | XP_041432378.1 | 2371 | germinal-center associated nuclear protein | |
| 397843 | mcm4.L | NM_001087979.1 | 3051 | NP_001081448.1 | 863 | DNA replication licensing factor mcm4-B | |
| 373601 | mcm4.S | NM_001085600.2 | 3055 | NP_001079069.1 | 858 | DNA replication licensing factor mcm4-A | |
| 380587 | mcm5.L | NM_001087424.1 | 2994 | NP_001080893.1 | 735 | DNA replication licensing factor mcm5-A | |
| 379699 | mcm5.S | NM_001086540.1 | 3042 | NP_001080009.1 | 735 | DNA replication licensing factor mcm5-B | |
| 398071 | mcm6.2.L | NM_001088353.1 | 2642 | NP_001081822.1 | 821 | maternal DNA replication licensing factor mcm6 | |
| 108705347 | mcm6.2.S | XM_018242191.2 | 759 | XP_018097680.2 | 168 | maternal DNA replication licensing factor mcm6-like | |
| 394426 | mcm6.L | NM_001137567.1 | 2790 | NP_001131039.1 | 823 | zygotic DNA replication licensing factor mcm6-A | |
| 380282 | mcm6.S | NM_001087121.1 | 3422 | NP_001080590.1 | 825 | zygotic DNA replication licensing factor mcm6-B | |
| 380414 | mcm7.L | NM_001087253.1 | 2557 | NP_001080722.1 | 720 | DNA replication licensing factor mcm7-B | |
| 397852 | mcm7.S | NM_001087997.1 | 2547 | NP_001081466.1 | 720 | DNA replication licensing factor mcm7-A | |
| 380250 | mcmbp.L | NM_001087089.1 | 2142 | NP_001080558.1 | 626 | mini-chromosome maintenance complex-binding protein | |
| 431919 | mtbp.S | NM_001091401.1 | 3024 | NP_001084870.1 | 860 | mdm2-binding protein | |
| 398063 | orc1.L | NM_001088337.1 | 3208 | NP_001081806.1 | 886 | origin recognition complex subunit 1 L homeolog | |
| 394365 | orc2.S | NM_001087601.1 | 2056 | NP_001081070.1 | 558 | origin recognition complex subunit 2 | |
| 379084 | orc3.S | NM_001085928.1 | 2301 | NP_001079397.1 | 709 | origin recognition complex subunit 3 S homeolog | |
| 397923 | orc4.L | NM_001088092.1 | 1444 | NP_001081561.1 | 432 | origin recognition complex subunit 4 | |
| 734494 | orc5.L | NM_001095975.1 | 1575 | NP_001089444.1 | 448 | origin recognition complex subunit 5 L homeolog | |
| 446747 | orc6.L | NM_001093443.1 | 2323 | NP_001086912.1 | 225 | origin recognition complex subunit 6 L homeolog | |
| 398425 | pcna.L | NM_001088895.1 | 1119 | NP_001082364.1 | 261 | proliferating cell nuclear antigen L homeolog | |
| 394328 | pcna.S | NM_001087542.1 | 1051 | NP_001081011.1 | 261 | proliferating cell nuclear antigen | |
| 108701628 | plk1.L | XM_018236516.2 | 2395 | XP_018092005.1 | 598 | serine/threonine-protein kinase PLK1-like | |
| 380481 | plk1.S | NM_001087319.1 | 2381 | NP_001080788.1 | 598 | serine/threonine-protein kinase PLK1 | |
| 398200 | pola1.S | NM_001088586.1 | 4629 | NP_001082055.1 | 1458 | DNA polymerase alpha catalytic subunit | |
| 446807 | pola2.L | NM_001093503.1 | 3160 | NP_001086972.1 | 598 | polymerase (DNA directed), alpha 2, accessory subunit L homeolog | |
| 447518 | pold1.L | NM_001094225.1 | 4106 | NP_001087694.1 | 1109 | DNA-directed DNA polymerase delta 1 | |
| 379793 | pold2.L | NM_001086632.1 | 1736 | NP_001080101.1 | 463 | DNA polymerase delta subunit 2 | |
| 398707 | pold3.L | NM_001089290.1 | 1801 | NP_001082759.1 | 454 | DNA-directed DNA polymerase delta 3 | |
| 394427 | pole.L | XM_018260369.2 | 7576 | XP_018115858.1 | 2285 | DNA polymerase epsilon catalytic subunit A | |
| 399116 | pole2.L | NM_001090314.1 | 1786 | NP_001083783.1 | 527 | DNA-directed DNA polymerase epsilon 2 | |
| 407748 | pole3.L | XM_018228794.2 | 1295 | XP_018084283.1 | 147 | DNA polymerase epsilon subunit 3 | |
| 779084 | pole4.L | NM_001096724.1 | 360 | NP_001090193.1 | 116 | DNA-directed DNA polymerase epsilon 4 | |
| 397767 | ppp1cc.L | NM_001087839.1 | 2065 | NP_001081308.1 | 323 | serine/threonine-protein phosphatase PP1-gamma catalytic subunit A | |
| 380598 | ppp1cc.S | NM_001087435.1 | 1926 | NP_001080904.1 | 323 | serine/threonine-protein phosphatase PP1-gamma catalytic subunit B | |
| 443852 | ppp1cb.L | NM_001091957.1 | 1171 | NP_001085426.1 | 327 | serine/threonine-protein phosphatase PP1-beta catalytic subunit | |
| 379914 | ppp1ca.L | NM_001086753.1 | 1908 | NP_001080222.1 | 326 | protein phosphatase 1, catalytic subunit, alpha isozyme L homeolog | |
| 108704179 | ppp1ca.S | XM_041578147.1 | 2000 | XP_041434081.1 | 326 | serine/threonine-protein phosphatase alpha-2 | |
| 108717671 | ppp1cb.S | XM_018264908.2 | 1845 | XP_018120397.1 | 327 | serine/threonine-protein phosphatase PP1-beta catalytic subunit | |
| 733317 | recql4.L | NM_001095632.2 | 4727 | NP_001089101.2 | 1504 | RECQL4-helicase-like protein | |
| 100380972 | rfc1.L | XM_018229167.2 | 3948 | XP_018084656.1 | 1147 | replication factor C subunit 1 | |
| 431883 | rfc2.L | NM_001091368.1 | 1224 | NP_001084837.1 | 348 | replication factor C subunit 2 L homeolog | |
| 108709658 | rfc2.S | XM_041584126.1 | 1217 | XP_041440060.1 | 255 | replication factor C subunit 2-like | |
| 734626 | rfc3.L | NM_001096101.1 | 1885 | NP_001089570.1 | 356 | replication factor C subunit 3 L homeolog | |
| 398706 | rfc4.L | NM_001089288.1 | 1256 | NP_001082757.1 | 363 | replication factor C subunit 4 L homeolog | |
| 380369 | rfc5.L | NM_001087208.1 | 1306 | NP_001080677.1 | 335 | replication factor C subunit 5 L homeolog | |
| 443952 | rfc5.S | NM_001092057.1 | 1382 | NP_001085526.1 | 335 | replication factor C subunit 5 S homeolog | |
| 101027299 | rif1.S | NM_001280649.1 | 6984 | NP_001267578.1 | 2327 | replication timing regulatory factor 1 S homeolog | |
| 397937 | rpa1.L | NM_001088116.1 | 2056 | NP_001081585.1 | 609 | replication protein A 70 kDa DNA-binding subunit | |
| 108708846 | rpa1.S | XM_041583312.1 | 2498 | XP_041439246.1 | 611 | replication protein A 70 kDa DNA-binding subunit-like | |
| 443819 | rpa2.L | NM_001091924.1 | 1310 | NP_001085393.1 | 276 | replication protein A 32 kDa subunit-A | |
| 100036853 | rpa2.S | NM_001097140.1 | 1384 | NP_001090609.1 | 274 | replication protein A 32 kDa subunit-B | |
| 495401 | rpa3.L | NM_001095060.1 | 891 | NP_001088529.1 | 121 | replication protein A3 L homeolog | |
| 446650 | rpa3.S | NM_001093346.1 | 868 | NP_001086815.1 | 121 | replication protein A3 S homeolog | |
| 108712846 | ticrr.S | XM_018255324.2 | 6860 | XP_018110813.1 | 2020 | treslin-like | isoform X1 |
| 398573 | topbp1.L | NM_001089099.1 | 5078 | NP_001082568.1 | 1513 | DNA topoisomerase II binding protein 1 L homeolog | |
| 100653494 | yap1.L | XM_041582680.1 | 2235 | XP_041438614.1 | 459 | transcriptional coactivator YAP1-A | isoform X1 |
| 100381098 | yap1.S | NM_001174024.1 | 2679 | NP_001167495.1 | 335 | transcriptional coactivator YAP1-B | |
| 379850 | zmcm3.L | NM_001086689.1 | 3143 | NP_001080158.1 | 806 | zygotic DNA replication licensing factor mcm3 | |

### Supplementary Table 5: Replication genes *X. eysoole*

| Gene name | X. laevis protein | Gene | Chromosome | X.laevis | X. eysoole | X.laevis transcript length | X.eysoole transcript length | Orthologs |
|---|---|---|---|---|---|---|---|---|
| ATM serine/threonine kinase | atm.L | atm | L | NP 001081968 | TRINITY GG 87 c2649 g10 i1 | 9186 | 0 | 1 |
| ATR serine/threonine kinase | atr.L | atr | L | NP 001082049 | TRINITY GG 20 c913 g1 i1 | 8346 | 0 | 1 |
| cell division cycle 45 | cdc45.L | cdc45 | S | XP 018093132 | TRINITY GG 94 c6265 g1 i3 | 2079 | 1985 | 1 |
| cell division cycle 6 | cdc6.L | cdc6 | L | NP 001084440 | TRINITY GG 44 c176 g1 i2 | 2213 | 2315 | 1 |
| cell division cycle 6 | cdc6.S | cdc6 | S | NP 001081844 | TRINITY GG 70 c8140 g1 i1 | 2323 | 2231 | 1 |
| cell division cycle 7 | cdc7.L | cdc7 | L | NP 001081878 | TRINITY GG 51 c889 g1 i10 | 1665 | 1594 | 1 |
| cyclin-dependent kinase 2 | cdk2.L | cdk2 | L | NP 001084976 | TRINITY GG 87 c4153 g1 i1 | 2391 | 1637 | 1 |
| cyclin-dependent kinase 2 | cdk2.S | cdk2 | S | NP 001084120 | TRINITY GG 60 c339 g1 i1 | 919 | 2007 | 1 |
| cyclin-dependent kinase 2 associated protein 1 | cdk2ap1.L | cdk2ap1 | L | XP 018118303 | ND | 1424 | ND | 0 |
| cyclin-dependent kinase 2 associated protein 1 | cdk2ap1.S | cdk2ap1 | S | XP 041436285 | TRINITY GG 94 c781 g1 i4 | 1640 | 1443 | 1 |
| chromatin licensing and DNA replication factor 1 | cdt1.L | cdt1 | L | NP 001081738 | TRINITY GG 51 c1390 g1 i6 | 2912 | 2615 | 1 |
| chromatin licensing and DNA replication factor 1 | cdt1.S | cdt1 | S | XP 018115992 | TRINITY GG 9 c283 g1 i4 | 3681 | 2072 | 1 |
| DBF4 zinc finger | dbf4.L | dbf4 | L | XP 018121834 | TRINITY GG 77 c149 g1 i3 | 2323 | 2539 | 1 |
| DBF4 zinc finger B | dbf4b.L | dbf4b | L | NP 001082740 | TRINITY GG 44 c1196 g1 i1 | 3811 | 0 | 0 |
| downstream neighbor of SON | donson.L | donson | L | XP 041437004 | TRINITY GG 86 c404 g1 i7 | 2651 | 1924 | 1 |
| flap structure-specific endonuclease 1 | fen1.S | fen1 | L | NP 001080960 | TRINITY GG 50 c374 g1 i4 | 1357 | 1488 | 1 |
| flap structure-specific endonuclease 1 | fen1.L | fen1 | S | NP 001080984 | TRINITY GG 9 c267 g1 i2 | 1469 | 1507 | 1 |
| GINS complex subunit 1 (Psf1 homolog) | gins1.S | gins1 | S | NP 001085502 | TRINITY GG 4 c800 g1 i1 | 1056 | 870 | 1 |
| GINS complex subunit 2 (Psf2 homolog) | gins2.L | gins2 | L | NP 001085627 | TRINITY GG 51 c210 g1 i9 | 1084 | 1346 | 1 |
| GINS complex subunit 2 (Psf2 homolog) | gins4.L | gins4 | L | XP 018106356 | TRINITY GG 72 c27 g1 i20 | 1150 | 1715 | 1 |
| geminin, DNA replication inhibitor L homeolog | gmnn.L | gmnn | L | NP 001081872 | TRINITY GG 78 c128 g1 i8 | 1047 | 1063 | 1 |
| geminin, DNA replication inhibitor S homeolog | gmnn.S | gmnn | S | NP 001084216 | ND | 1096 | ND | 0 |
| DNA ligase 1 | lig1.L | lig1 | L | NP 001081653 | TRINITY GG 35 c57 g2 i1 | 3770 | 3776 | 1 |
| minichromosome maintenance 10 replication initiation factor | mcm10.L | mcm10 | L | XP 018106300 | TRINITY GG 73 c163 g3 i2 | 3510 | 3530 | 1 |
| minichromosome maintenance complex component 2 | mcm2.L | mcm2 | L | NP 001080759 | TRINITY GG 51 c26 g1 i2 | 3278 | 3360 | 1 |
| minichromosome maintenance complex component 2 | mcm2.S | mcm2 | S | XP 018116882 | TRINITY GG 10 c315 g1 i1 | 3644 | 3923 | 1 |
| minichromosome maintenance complex component 3 | mcm3.L | mcm3 | L | NP 001081412 | ND | 2548 | ND | 0 |
| minichromosome maintenance complex component 3 associated protein | mcm3ap.L | mcm3ap | L | XP 018091512 | TRINITY GG 43 c81 g1 i13 | 8433 | 9174 | 1 |
| minichromosome maintenance complex component 4 | mcm4.L | mcm4 | L | NP 001081448 | TRINITY GG 78 c97 g1 i4 | 3051 | 3230 | 1 |
| minichromosome maintenance complex component 4 | mcm4.S | mcm4 | S | NP 001079069 | ND | 2548 | ND | 0 |
| minichromosome maintenance complex component 5 | mcm5.L | mcm5 | L | NP 001080893 | TRINITY GG 51 c364 g1 i2 | 2994 | 3133 | 1 |
| minichromosome maintenance complex component 5 | mcm5.S | mcm5 | S | NP 001080009 | TRINITY GG 10 c147 g1 i2 | 3042 | 3915 | 1 |
| minichromosome maintenance complex component 6 | mcm6.L | mcm6 | L | NP 001131039 | TRINITY GG 43 c41 g1 i3 | 2790 | 3231 | 1 |
| minichromosome maintenance complex component 7 | mcm6.S | mcm6 | S | NP 001080590 | TRINITY GG 70 c58 g1 i5 | 3422 | 3103 | 1 |
| minichromosome maintenance complex component 6 gene 2 | mcm6.2.L | mcm6.2 | L | NP 001131039 | ND | 2790 | ND | 0 |
| minichromosome maintenance complex component 6 gene 2 | mcm6.2.S | mcm6.2 | S | NP 001080590 | ND | 3422 | ND | 0 |
| minichromosome maintenance complex component 7 | mcm7.L | mcm7 | L | NP 001080722 | TRINITY GG 72 c116 g1 i19 | 2557 | 2579 | 1 |
| minichromosome maintenance complex component 7 | mcm7.S | mcm7 | S | NP 001081466 | TRINITY GG 11 c2882 g1 i1 | 2547 | 2589 | 1 |
| minichromosome maintenance complex binding protein | mcmbp.L | mcmbp | L | NP 001080558 | TRINITY GG 37 c974 g1 i6 | 2142 | 2286 | 1 |
| MDM2 binding protein | mtbp.S | mtbp | S | NP 001084870 | ND | 3024 | ND | 0 |
| origin recognition complex subunit 1 | orc1.L | orc1 | L | NP 001081806 | TRINITY GG 51 c930 g1 i4 | 3208 | 2955 | 1 |
| origin recognition complex subunit 2 | orc2.S | orc2 | S | XP 018092774 | TRINITY GG 70 c1105 g1 i3 | 2139 | 2172 | 1 |
| origin recognition complex subunit 3 | orc3.S | orc3 | S | XP 018119715 | TRINITY GG 3 c4093 g1 i1 | 2722 | 2346 | 1 |
| origin recognition complex subunit 4 | orc4.L | orc4 | L | NP 001081561 | TRINITY GG 43 c223 g1 i6 | 1444 | 1514 | 1 |
| origin recognition complex subunit 5 | orc5.L | orc5 | L | NP 001089444 | TRINITY GG 78 c823 g1 i6 | 1575 | 1812 | 1 |
| origin recognition complex subunit 6 | orc6.L | orc6 | L | NP 001086912 | TRINITY GG 51 c473 g1 i6 | 2323 | 1944 | 1 |
| proliferating cell nuclear antigen | pcna.S | pcna | L | NP 001082364 | TRINITY GG 72 c1301 g1 i1 | 1119 | 1267 | 1 |
| proliferating cell nuclear antigen | pcna.L | pcna | S | NP 001081011 | TRINITY GG 13 c116 g1 i1 | 1051 | 1158 | 1 |
| polo like kinase 1 | plk1.L | plk1 | L | XP 018092005 | TRINITY GG 49 c568 g1 i8 | 2395 | 2395 | 1 |
| polo like kinase 1 | plk1.S | plk1 | S | NP 001080788 | TRINITY GG 69 c53 g1 i5 | 2381 | 2382 | 1 |
| polymerase (DNA directed), alpha 1, catalytic subunit | pola1.S | pola1 | S | NP 001082055 | TRINITY GG 54 c103 g1 i2 | 4629 | 5058 | 1 |
| polymerase (DNA directed), alpha 2, accessory subunit | pola2.L | pola2 | L | XP 018112289 | TRINITY GG 50 c2973 g1 i4 | 2505 | 2550 | 1 |
| polymerase (DNA directed), delta 1, catalytic subunit | pold1.L | pold1 | L | NP 001087694 | TRINITY GG 40 c131 g1 i8 | 4106 | 4117 | 1 |
| polymerase (DNA directed), delta 2, accessory subunit | pold2.L | pold2 | L | XP 018106174 | TRINITY GG 72 c201 g1 i3 | 1755 | 1712 | 1 |
| polymerase (DNA-directed), delta 3, accessory subunit | pold3.L | pold3 | L | NP 001082759 | TRINITY GG 87 c289 g1 i10 | 1801 | 1766 | 1 |
| polymerase (DNA directed), epsilon, catalytic subunit A | pole.L | pole | L | XP 018115858 | ND | 7576 | ND | 0 |
| polymerase (DNA directed), epsilon 2, catalytic subunit | pole2.L | pole2 | L | NP 001083783 | TRINITY GG 17 c1857 g1 i6 | 1786 | 3215 | 1 |
| polymerase (DNA directed), epsilon 3, catalytic subunit | pole3.L | pole3 | L | XP 018084282 | TRINITY GG 16 c199 g1 i6 | 1250 | 1392 | 1 |
| polymerase (DNA directed), epsilon 4, catalytic subunit | pole4.L | pole4 | L | NP 001090193 | TRINITY GG 73 c4609 g1 i1 | 360 | 860 | 1 |
| protein phosphatase 1, catalytic subunit, alpha isozyme L homeolog | ppp1ca.L | ppp1ca | L | NP 001080222 | TRINITY GG 48 c52 g1 i3 | 1908 | 2013 | 1 |
| serine/threonine-protein phosphatase alpha-2 | ppp1ca.S | ppp1ca | S | XP 041434081 | TRINITY GG 68 c39 g1 i1 | 2000 | 1932 | 1 |
| serine/threonine-protein phosphatase PP1-beta catalytic subunit | ppp1cb.L | ppp1cb | L | NP 001085426 | TRINITY GG 21 c747 g1 i5 | 1171 | 2590 | 1 |
| serine/threonine-protein phosphatase PP1-beta catalytic subunit | ppp1cb.S | ppp1cb | S | XP 018120397 | TRINITY GG 4 c47 g1 i4 | 1845 | 1307 | 1 |
| serine/threonine-protein phosphatase PP1-gamma catalytic subunit A | ppp1cc.L | ppp1cc | L | NP 001081308 | ND | 2065 | ND | 0 |
| serine/threonine-protein phosphatase PP1-gamma catalytic subunit B | ppp1cc.S | ppp1cc | S | NP 001080904 | TRINITY GG 94 c109 g2 i3 | 1926 | 1931 | 1 |
| RecQ helicase-like 4 | recql4.L | recql4 | L | NP 001089101 | TRINITY GG 86 c463 g1 i2 | 4727 | 3659 | 1 |
| replication factor C subunit1 | rfc1.L | rfc1 | L | XP 018084656 | TRINITY GG 24 c253 g1 i4 | 3948 | 3514 | 1 |
| replication factor C subunit2 | rfc2.S | rfc2 | S | XP 041440060 | TRINITY GG 53 c149 g1 i1 | 1217 | 1031 | 1 |
| replication factor C subunit2 | rfc2.L | rfc2 | L | NP 001084837 | TRINITY GG 87 c240 g1 i6 | 1224 | 1414 | 2 |
| replication factor C subunit3 | rfc3.L | rfc3 | L | NP 001089570 | TRINITY GG 87 c1728 g1 i5 | 1885 | 2471 | 1 |
| replication factor C subunit4 | rfc4.L | rfc4 | L | NP 001082757 | TRINITY GG 20 c735 g1 i4 | 1256 | 1390 | 1 |
| replication factor C subunit5 | rfc5.L | rfc5 | L | NP 001080677 | ND | 1306 | ND | 0 |
| replication factor C subunit5 | rfc5.S | rfc5 | S | NP 001085526 | TRINITY GG 94 c389 g1 i1 | 1382 | 1561 | 1 |
| replication timing regulatory factor 1, 2 isoforms X1, X2 | rif1.S | rif1 | S | NP 001267578 | TRINITY GG 70 c107 g1 i5 | 6984 | 8327 | 1 |
| replication protein A1 | rpa1.L | rpa1 | L | NP 001081585 | TRINITY GG 86 c275 g1 i2 | 2056 | 3567 | 1 |
| replication protein A1 | rpa1.S | rpa1 | S | XP 041439246 | TRINITY GG 54 c1677 g1 i1 | 2498 | 1831 | 1 |
| replication protein A2 | rpa2.S | rpa2 | S | NP 001090609 | TRINITY GG 59 c378 g1 i1 | 1384 | 783 | 1 |
| replication protein A2 | rpa2.L | rpa2 | L | NP 001085393 | TRINITY GG 86 c369 g1 i1 | 1310 | 1750 | 1 |
| replication protein A3 | rpa3.L | rpa3 | L | NP 001088529 | TRINITY GG 77 c471 g1 i5 | 891 | 873 | 1 |
| replication protein A3 | rpa3.S | rpa3 | S | NP 001086815 | TRINITY GG 80 c1093 g1 i14 | 868 | 921 | 1 |
| TOPBP1-interacting checkpoint and replication regulator | ticrr | ticrr | S | XP 018110813 | TRINITY GG 72 c316 g1 i1 | 6860 | 5764 | 1 |
| DNA topoisomerase II binding protein 1 | topbp1 | topbp1 | L | NP 001082568 | TRINITY GG 77 c421 g1 i1 | 5078 | 5035 | 1 |
| Yes associated protein 1 | yap1.L | yap1 | L | XP 041438614 | TRINITY GG 87 c1521 g1 i1 | 2235 | 2402 | 1 |
| Yes associated protein 1 | yap1.S | yap1 | S | NP 001167495 | TRINITY GG 60 c5527 g1 i3 | 2679 | 1274 | 1 |
| Putative ortholog of DNA replication licensing factor MCM3 | zmcm3.L | zmcm3 | L | NP 001080158 | TRINITY GG 19 c18 g1 i5 | 3143 | 3523 | 1 |
| | | | | | | | 11 ND | 0,870588235 |

### Supplementary Table 6: Comparison of conservation of replication genes *X. tropicalis/X. laevis*

| gene | subgenome | X. laevis gene | qseqid | sseqid | qlen (aa) | slen (nt) | pident | length (aa) | mismatch | gapopen | qstart | qend | sstart | send | qcov |
|---|---|---|---|---|---|---|---|---|---|---|---|---|---|---|---|
| atm | L | atm.L | NP_001081968.1 | XM_002934928.5 | 3061 | 10039 | 91,5 | 3062 | 257 | 3 | 1 | 3061 | 152 | 9331 | 1,00 |
| atr | L | atr.L | NP_001082049.1 | XM_031902384.1 | 2655 | 8400 | 93,6 | 2656 | 168 | 2 | 1 | 2655 | 48 | 8012 | 1,00 |
| cdc45 | S | cdc45.S | NP_001087102.1 | XM_002931882.5 | 567 | 2027 | 95,3 | 569 | 25 | 1 | 1 | 567 | 115 | 1821 | 1,00 |
| cdc6 | L | cdc6.L | NP_001084440.1 | NM_001007993.1 | 554 | 2232 | 89,8 | 558 | 49 | 3 | 1 | 554 | 77 | 1738 | 1,01 |
| cdc6 | S | cdc6.S | NP_001081844.1 | NM_001007993.1 | 554 | 2232 | 89,7 | 555 | 55 | 2 | 1 | 554 | 77 | 1738 | 1,00 |
| cdc7 | L | cdc7.L | NP_001081878.1 | NM_001016581.2 | 483 | 1687 | 90,9 | 485 | 40 | 2 | 1 | 483 | 114 | 1562 | 1,00 |
| cdk2 | L | cdk2.L | NP_001084976.1 | NM_001008135.1 | 297 | 1903 | 97,6 | 297 | 7 | 0 | 1 | 297 | 77 | 967 | 1,00 |
| cdk2 | S | cdk2.S | NP_001084120.1 | NM_001008135.1 | 297 | 1903 | 97,6 | 297 | 7 | 0 | 1 | 297 | 77 | 967 | 1,00 |
| cdk2ap1 | L | cdk2ap1.L | XP_018118294.1 | NM_001016730.2 | 120 | 1238 | 98,3 | 120 | 2 | 0 | 1 | 120 | 9 | 368 | 1,00 |
| cdk2ap1 | S | cdk2ap1.S | XP_041436285.1 | NM_001016730.2 | 120 | 1238 | 95,8 | 120 | 5 | 0 | 1 | 120 | 9 | 368 | 1,00 |
| cdt1 | L | cdt1.L | NP_001081738.1 | XM_002933654.5 | 620 | 2717 | 88,1 | 621 | 69 | 3 | 1 | 620 | 145 | 1995 | 1,00 |
| cdt1 | S | cdt1.S | XP_018115992.1 | XM_002933654.5 | 618 | 2717 | 88,1 | 623 | 63 | 6 | 1 | 618 | 145 | 1995 | 1,01 |
| dbf4 | L | dbf4.L | NP_001082606.1 | NM_001044505.1 | 661 | 2493 | 79,0 | 668 | 128 | 4 | 1 | 661 | 156 | 2144 | 1,01 |
| dbf4b | L | dbf4b.L | NP_001082740.1 | XM_004918590.4 | 784 | 4123 | 76,3 | 801 | 166 | 9 | 5 | 783 | 128 | 2524 | 1,02 |
| donson | L | donson.L | NP_001088556.1 | NM_204053.1 | 579 | 2399 | 89,5 | 581 | 54 | 3 | 1 | 579 | 57 | 1784 | 1,00 |
| fen1 | L | fen1.L | NP_001080960.1 | NM_001017005.2 | 382 | 1436 | 94,3 | 353 | 20 | 0 | 1 | 353 | 79 | 1137 | 0,92 |
| fen1 | S | fen1.S | NP_001080984.1 | NM_001017005.2 | 382 | 1436 | 94,8 | 382 | 20 | 0 | 1 | 382 | 79 | 1224 | 1,00 |
| gins1 | S | gins1.S | NP_001085502.1 | NM_001016164.2 | 196 | 752 | 97,4 | 196 | 5 | 0 | 1 | 196 | 25 | 612 | 1,00 |
| gins2 | L | gins2.L | NP_001085627.1 | NM_001017015.2 | 185 | 1483 | 96,8 | 185 | 6 | 0 | 1 | 185 | 2 | 556 | 1,00 |
| gins4 | L | gins4.L | NP_001084702.1 | NM_001011045.1 | 221 | 1744 | 96,4 | 221 | 8 | 0 | 1 | 221 | 106 | 768 | 1,00 |
| gmnn | L | gmnn.L | NP_001081872.1 | NM_001039736.1 | 216 | 1047 | 86,8 | 219 | 26 | 1 | 1 | 216 | 28 | 684 | 1,01 |
| gmnn | S | gmnn.S | NP_001084216.1 | NM_001039736.1 | 219 | 1047 | 90,9 | 219 | 20 | 0 | 1 | 219 | 28 | 684 | 1,00 |
| lig1 | L | lig1.L | NP_001081653.1 | XM_002941883.5 | 1070 | 4372 | 85,3 | 1073 | 122 | 5 | 1 | 1070 | 156 | 3275 | 1,00 |
| mcm10 | L | mcm10.L | NP_001082047.1 | NM_001016909.2 | 860 | 3072 | 86,6 | 837 | 95 | 5 | 23 | 858 | 151 | 2613 | 0,97 |
| mcm2 | L | mcm2.L | NP_001080759.1 | NM_001006771.1 | 886 | 3268 | 96,7 | 886 | 27 | 1 | 1 | 886 | 54 | 2705 | 1,00 |
| mcm2 | S | mcm2.S | XP_018116882.1 | NM_001006771.1 | 885 | 3268 | 96,4 | 886 | 29 | 2 | 1 | 885 | 54 | 2705 | 1,00 |
| mcm3 | L | mcm3.L | NP_001081412.1 | XM_002934119.5 | 807 | 2578 | 96,3 | 806 | 30 | 0 | 1 | 806 | 39 | 2456 | 1,00 |
| mcm3ap | L | mcm3ap.L | XP_041432378.1 | XM_004917891.2 | 2371 | 8220 | 80,5 | 2395 | 420 | 15 | 1 | 2370 | 113 | 7234 | 1,01 |
| mcm4 | L | mcm4.L | NP_001081448.1 | NM_001005655.1 | 863 | 3290 | 97,4 | 842 | 22 | 0 | 22 | 863 | 162 | 2687 | 0,98 |
| mcm4 | S | mcm4.S | NP_001079069.1 | NM_001005655.1 | 858 | 3290 | 97,7 | 795 | 18 | 0 | 64 | 858 | 303 | 2687 | 0,93 |
| mcm5 | L | mcm5.L | NP_001080893.1 | NM_001017327.3 | 735 | 3028 | 98,5 | 735 | 11 | 0 | 1 | 735 | 16 | 2220 | 1,00 |
| mcm5 | S | mcm5.S | NP_001080009.1 | NM_001017327.3 | 735 | 3028 | 98,0 | 735 | 15 | 0 | 1 | 735 | 16 | 2220 | 1,00 |
| mcm6 | L | mcm6.L | NP_001131039.1 | NM_204062.1 | 823 | 2893 | 94,9 | 823 | 42 | 0 | 1 | 823 | 49 | 2517 | 1,00 |
| mcm6 | S | mcm6.S | NP_001080590.1 | NM_204062.1 | 825 | 2893 | 95,4 | 826 | 34 | 2 | 1 | 825 | 49 | 2517 | 1,00 |
| mcm6.2 | L | mcm6.2.L | NP_001081822.1 | NM_001016221.2 | 821 | 2625 | 96,5 | 822 | 27 | 2 | 1 | 821 | 20 | 2482 | 1,00 |
| mcm6.2 | S | mcm6.2.S | XP_018097680.2 | NM_001016221.2 | 168 | 2625 | 86,7 | 166 | 19 | 3 | 1 | 165 | 1451 | 1942 | 0,99 |
| mcm7 | L | mcm7.L | NP_001080722.1 | NM_213712.1 | 720 | 2499 | 97,1 | 720 | 21 | 0 | 1 | 720 | 34 | 2193 | 1,00 |
| mcm7 | S | mcm7.S | NP_001081466.1 | NM_213712.1 | 720 | 2499 | 97,8 | 720 | 16 | 0 | 1 | 720 | 34 | 2193 | 1,00 |
| mcmbp | L | mcmbp.L | NP_001080558.1 | NM_001008082.1 | 626 | 2250 | 87,6 | 627 | 77 | 1 | 1 | 626 | 121 | 2001 | 1,00 |
| mtbp | S | mtbp.S | NP_001084870.1 | NM_001114218.1 | 860 | 3195 | 88,8 | 866 | 90 | 3 | 1 | 859 | 66 | 2663 | 1,01 |
| orc1 | L | orc1.L | NP_001081806.1 | NM_001045729.1 | 886 | 2859 | 90,7 | 889 | 79 | 3 | 1 | 886 | 29 | 2692 | 1,00 |
| orc2 | S | orc2.S | NP_001081070.1 | NM_001079338.1 | 558 | 2016 | 92,3 | 558 | 43 | 0 | 1 | 558 | 28 | 1701 | 1,00 |
| orc3 | S | orc3.S | NP_001079397.1 | NM_001103067.1 | 709 | 2257 | 91,5 | 709 | 60 | 0 | 1 | 709 | 7 | 2133 | 1,00 |
| orc4 | L | orc4.L | NP_001081561.1 | NM_001005654.1 | 432 | 1557 | 93,7 | 428 | 27 | 0 | 1 | 428 | 110 | 1393 | 0,99 |
| orc5 | L | orc5.L | NP_001089444.1 | NM_001017013.2 | 448 | 1815 | 94,4 | 448 | 25 | 0 | 1 | 448 | 150 | 1493 | 1,00 |
| orc6 | L | orc6.L | NP_001086912.1 | NM_001017185.2 | 225 | 1148 | 93,4 | 212 | 14 | 0 | 1 | 212 | 190 | 825 | 0,94 |
| pcna | L | pcna.L | NP_001082364.1 | NM_001007920.1 | 261 | 1134 | 99,2 | 261 | 2 | 0 | 1 | 261 | 118 | 900 | 1,00 |
| pcna | S | pcna.S | NP_001081011.1 | NM_001007920.1 | 261 | 1134 | 99,2 | 261 | 2 | 0 | 1 | 261 | 118 | 900 | 1,00 |
| plk1 | L | plk1.L | XP_018092005.1 | NM_213679.2 | 598 | 2178 | 96,7 | 598 | 20 | 0 | 1 | 598 | 147 | 1940 | 1,00 |
| plk1 | S | plk1.S | NP_001080788.1 | NM_213679.2 | 598 | 2178 | 96,6 | 585 | 20 | 0 | 1 | 585 | 147 | 1901 | 0,98 |
| pola1 | S | pola1.S | NP_001082055.1 | XM_031896702.1 | 1458 | 4536 | 92,3 | 1419 | 97 | 6 | 1 | 1411 | 106 | 4350 | 0,97 |
| pola2 | L | pola2.L | NP_001086972.1 | NM_001005060.1 | 598 | 2765 | 94,0 | 598 | 35 | 1 | 1 | 598 | 182 | 1972 | 1,00 |
| pold1 | L | pold1.L | NP_001087694.1 | NM_001045739.1 | 1109 | 3993 | 96,3 | 1109 | 41 | 0 | 1 | 1109 | 98 | 3424 | 1,00 |
| pold2 | L | pold2.L | NP_001080101.1 | NM_001005794.1 | 463 | 1701 | 95,0 | 463 | 23 | 0 | 1 | 463 | 75 | 1463 | 1,00 |
| pold3 | L | pold3.L | NP_001082759.1 | NM_001016556.2 | 454 | 1314 | 87,0 | 207 | 26 | 1 | 1 | 207 | 64 | 681 | 0,46 |
| pole | L | pole.L | XP_018115858.1 | XM_004910635.4 | 2285 | 7100 | 97,6 | 2286 | 55 | 1 | 1 | 2285 | 82 | 6939 | 1,00 |
| pole2 | L | pole2.L | NP_001083783.1 | NM_001030470.1 | 527 | 1718 | 96,8 | 527 | 17 | 0 | 1 | 527 | 26 | 1606 | 1,00 |
| pole3 | L | pole3.L | XP_018084283.1 | NM_001017264.2 | 147 | 829 | 99,0 | 98 | 1 | 0 | 1 | 98 | 60 | 353 | 0,67 |
| pole4 | L | pole4.L | NP_001090193.1 | NM_001016605.2 | 116 | 1085 | 89,7 | 116 | 11 | 1 | 1 | 116 | 23 | 367 | 1,00 |
| ppp1ca | L | ppp1ca.L | NP_001080222 | NM_001127010 | 326 | 2141 | 100,0 | 326 | 0 | 0 | 1 | 326 | 71 | 1048 | 1,00 |
| ppp1ca | S | ppp1ca.S | XP_041434081 | NM_001127010 | 326 | 2141 | 99,4 | 326 | 2 | 0 | 1 | 326 | 71 | 1048 | 1,00 |
| ppp1cb | L | ppp1cb.L | NP_001085426 | NM_001011467 | 327 | 1183 | 100,0 | 327 | 0 | 0 | 1 | 327 | 7 | 987 | 1,00 |
| ppp1cb | S | ppp1cb.S | XP_018120397 | NM_001011467 | 327 | 1183 | 100,0 | 327 | 0 | 0 | 1 | 327 | 7 | 987 | 1,00 |
| ppp1cc | L | ppp1cc.L | NP_001081308 | NM_213670 | 323 | 1993 | 100,0 | 323 | 0 | 0 | 1 | 323 | 179 | 1147 | 1,00 |
| ppp1cc | S | ppp1cc.S | NP_001080904 | NM_213670 | 323 | 1993 | 99,7 | 323 | 1 | 0 | 1 | 323 | 179 | 1147 | 1,00 |
| recql4 | L | recql4.L | NP_001089101.2 | NM_001079414.1 | 1504 | 5033 | 80,8 | 1539 | 241 | 15 | 1 | 1504 | 41 | 4600 | 1,02 |
| rfc1 | L | rfc1.L | XP_018084656.1 | NM_001199567.1 | 1147 | 3988 | 89,4 | 1077 | 101 | 4 | 1 | 1074 | 185 | 3385 | 0,94 |
| rfc2 | L | rfc2.L | NP_001084837.1 | NM_001142016.1 | 348 | 1182 | 96,1 | 334 | 13 | 0 | 1 | 334 | 58 | 1059 | 0,96 |
| rfc2 | S | rfc2.S | XP_041440060.1 | NM_001142016.1 | 255 | 1182 | 85,6 | 229 | 32 | 1 | 1 | 228 | 55 | 741 | 0,90 |
| rfc3 | L | rfc3.L | NP_001089570.1 | XM_002934033.5 | 356 | 1802 | 96,6 | 356 | 12 | 0 | 1 | 356 | 146 | 1213 | 1,00 |
| rfc4 | L | rfc4.L | NP_001082757.1 | NM_001016363.2 | 363 | 1214 | 95,6 | 363 | 16 | 0 | 1 | 363 | 25 | 1113 | 1,00 |
| rfc5 | L | rfc5.L | NP_001080677.1 | NM_001011112.2 | 335 | 1895 | 97,3 | 335 | 9 | 0 | 1 | 335 | 107 | 1111 | 1,00 |
| rfc5 | S | rfc5.S | NP_001085526.1 | NM_001011112.2 | 335 | 1895 | 95,8 | 335 | 14 | 0 | 1 | 335 | 107 | 1111 | 1,00 |
| rif1 | S | rif1.S | NP_001267578.1 | XM_004917657.4 | 2327 | 8503 | 81,1 | 2379 | 378 | 22 | 1 | 2327 | 148 | 7224 | 1,02 |
| rpa1 | L | rpa1.L | NP_001081585.1 | NM_001015732.1 | 609 | 2979 | 94,1 | 610 | 34 | 2 | 1 | 609 | 111 | 1937 | 1,00 |
| rpa1 | S | rpa1.S | XP_041439246.1 | NM_001015732.1 | 611 | 2979 | 90,8 | 611 | 54 | 2 | 1 | 611 | 111 | 1937 | 1,00 |
| rpa2 | L | rpa2.L | NP_001085393.1 | NM_001006794.1 | 276 | 1221 | 94,7 | 245 | 13 | 0 | 32 | 276 | 118 | 852 | 0,89 |
| rpa2 | S | rpa2.S | NP_001090609.1 | NM_001006794.1 | 274 | 1221 | 88,7 | 247 | 26 | 1 | 28 | 274 | 118 | 852 | 0,90 |
| rpa3 | L | rpa3.L | NP_001088529.1 | NM_001016626.2 | 121 | 856 | 93,4 | 121 | 8 | 0 | 1 | 121 | 6 | 368 | 1,00 |
| rpa3 | S | rpa3.S | NP_001086815.1 | NM_001016626.2 | 121 | 856 | 89,3 | 121 | 13 | 0 | 1 | 121 | 6 | 368 | 1,00 |
| ticrr | S | ticrr.S | XP_018110813.1 | XM_031899228.1 | 2020 | 8000 | 79,0 | 2040 | 335 | 16 | 1 | 2020 | 222 | 6122 | 1,01 |
| topbp1 | L | topbp1.L | NP_001082568.1 | XM_002937745.4 | 1513 | 5547 | 90,6 | 1517 | 136 | 3 | 1 | 1513 | 138 | 4682 | 1,00 |
| yap1 | L | yap1.L | XP_041438614.1 | NM_001195768.1 | 459 | 1371 | 92,2 | 435 | 30 | 3 | 27 | 459 | 70 | 1368 | 0,95 |
| yap1 | S | yap1.S | NP_001167495.1 | XM_012956607.3 | 335 | 7414 | 92,3 | 261 | 19 | 1 | 76 | 335 | 876 | 1658 | 0,78 |
| zmcm3 | L | zmcm3.L | NP_001080158.1 | NM_001045766.1 | 806 | 3121 | 95,2 | 805 | 38 | 1 | 3 | 806 | 73 | 2487 | 1,00 |

### Supplementary Table 7: Comparison of conservation of replication genes
#### *X. laevis/X.eysoole*

| gene | subgenome | X. laevis gene | qseqid | sseqid | qlen (aa) | slen (nt) | pident | length (aa) | mismatch | gapopen | qstart | qend | sstart | send | qcov |
|---|---|---|---|---|---|---|---|---|---|---|---|---|---|---|---|
| atm | L | atm.L | NP_001081968.1 | TRINITY_GG_87_c2649_g10_i1 | 3061 | 2001 | 99,09 | 549 | 5 | 0 | 2513 | 3061 | 2001 | 355 | 0,18 |
| atr | L | atr.L | NP_001082049.1 | TRINITY_GG_20_c913_g1_i1 | 2655 | 4127 | 98,67 | 1279 | 17 | 0 | 1377 | 2655 | 4125 | 289 | 0,48 |
| cdc45 | S | cdc45.S | XP_018093132.1 | TRINITY_GG_94_c6265_g1_i3 | 576 | 1985 | 94,28 | 577 | 24 | 2 | 1 | 576 | 1902 | 196 | 1,00 |
| cdc6 | L | cdc6.L | NP_001084440.1 | TRINITY_GG_44_c176_g1_i2 | 554 | 2315 | 96,21 | 554 | 20 | 1 | 1 | 554 | 145 | 1803 | 1,00 |
| cdc6 | S | cdc6.S | NP_001081844.1 | TRINITY_GG_70_c8140_g1_i1 | 554 | 2231 | 94,62 | 558 | 26 | 1 | 1 | 554 | 2161 | 488 | 1,01 |
| cdc7 | L | cdc7.L | NP_001081878.1 | TRINITY_GG_51_c889_g1_i10 | 483 | 1594 | 94,43 | 485 | 25 | 1 | 1 | 483 | 75 | 1529 | 1,00 |
| cdk2 | L | cdk2.L | NP_001084976.1 | TRINITY_GG_87_c4153_g1_i1 | 297 | 1637 | 98,65 | 297 | 4 | 0 | 1 | 297 | 1537 | 647 | 1,00 |
| cdk2 | S | cdk2.S | NP_001084120.1 | TRINITY_GG_60_c339_g1_i1 | 297 | 2007 | 98,99 | 297 | 3 | 0 | 1 | 297 | 58 | 948 | 1,00 |
| cdk2ap1 | L | cdk2ap1.L | XP_018118303 | ND | 0 | 0 | 0,00 | 0 | 0 | 0 | 0 | 0 | 0 | 0 | 0 |
| cdk2ap1 | S | cdk2ap1.S | XP_041436285.1 | TRINITY_GG_94_c781_g1_i4 | 120 | 1443 | 100,00 | 120 | 0 | 0 | 1 | 120 | 1231 | 872 | 1,00 |
| cdt1 | L | cdt1.L | NP_001081738.1 | TRINITY_GG_51_c1390_g1_i6 | 620 | 2615 | 94,47 | 597 | 32 | 1 | 25 | 620 | 2 | 1792 | 0,96 |
| cdt1 | S | cdt1.S | XP_018115992.1 | TRINITY_GG_9_c283_g1_i4 | 618 | 2072 | 94,68 | 620 | 31 | 2 | 1 | 618 | 48 | 1907 | 1,00 |
| dbf4 | L | dbf4.L | XP_018121834.1 | TRINITY_GG_77_c149_g1_i3 | 672 | 2539 | 94,79 | 672 | 30 | 2 | 5 | 672 | 2339 | 327 | 1,00 |
| dbf4b | L | dbf4b.L | NP_001082740.1 | TRINITY_GG_44_c1196_g1_i1 | 784 | 1252 | 92,31 | 416 | 32 | 0 | 15 | 430 | 1250 | 3 | 0,53 |
| donson | L | donson.L | XP_041437004.1 | TRINITY_GG_86_c404_g1_i7 | 579 | 1924 | 96,20 | 579 | 21 | 1 | 1 | 579 | 1845 | 112 | 1,00 |
| fen1 | L | fen1.L | NP_001080960.1 | TRINITY_GG_50_c374_g1_i4 | 382 | 1488 | 98,30 | 353 | 6 | 0 | 1 | 353 | 1321 | 263 | 0,92 |
| fen1 | S | fen1.S | NP_001080984.1 | TRINITY_GG_9_c267_g1_i2 | 382 | 1507 | 97,91 | 382 | 8 | 0 | 1 | 382 | 152 | 1297 | 1,00 |
| gins1 | S | gins1.S | NP_001085502.1 | TRINITY_GG_4_c800_g1_i1 | 196 | 870 | 100,00 | 196 | 0 | 0 | 1 | 196 | 91 | 678 | 1,00 |
| gins2 | L | gins2.L | NP_001085627.1 | TRINITY_GG_51_c210_g1_i9 | 185 | 1346 | 98,92 | 185 | 2 | 0 | 1 | 185 | 91 | 645 | 1,00 |
| gins4 | L | gins4.L | XP_018106356.1 | TRINITY_GG_72_c27_g1_i20 | 221 | 1715 | 98,19 | 221 | 4 | 0 | 1 | 221 | 136 | 798 | 1,00 |
| gmnn | L | gmnn.L | NP_001081872.1 | TRINITY_GG_78_c128_g1_i8 | 216 | 1063 | 90,74 | 216 | 20 | 0 | 1 | 216 | 1007 | 360 | 1,00 |
| gmnn | S | gmnn.S | NP_001084216 | ND | 0 | 0 | 0,00 | 0 | 0 | 0 | 0 | 0 | 0 | 0 | 0,00 |
| lig1 | L | lig1.L | NP_001081653.1 | TRINITY_GG_35_c57_g2_i1 | 1070 | 3776 | 91,527 | 1074,00 | 53 | 4 | 1 | 1070 | 3619 | 500 | 1,00 |
| mcm10 | L | mcm10.L | XP_018106300.1 | TRINITY_GG_73_c163_g3_i2 | 860 | 3530 | 96,40 | 860 | 28 | 1 | 1 | 860 | 81 | 2651 | 1,00 |
| mcm2 | L | mcm2.L | NP_001080759.1 | TRINITY_GG_51_c26_g1_i2 | 886 | 3360 | 99,20 | 879 | 7 | 0 | 8 | 886 | 118 | 2754 | 0,99 |
| mcm2 | S | mcm2.S | XP_018116882.1 | TRINITY_GG_10_c315_g1_i1 | 885 | 3923 | 98,76 | 885 | 11 | 0 | 1 | 885 | 75 | 2729 | 1,00 |
| mcm3 | L | mcm3.L | NP_001081412 | ND | 0 | 0 | 0,00 | 0 | 0 | 0 | 0 | 0 | 0 | 0 | 0 |
| mcm3ap | L | mcm3ap.L | XP_018091512.1 | TRINITY_GG_43_c81_g1_i13 | 2371 | 9174 | 95,29 | 2039 | 93 | 3 | 335 | 2371 | 1 | 6114 | 0,86 |
| mcm4 | L | mcm4.L | NP_001081448.1 | TRINITY_GG_78_c97_g1_i4 | 863 | 3230 | 99,17 | 842 | 7 | 0 | 22 | 863 | 3081 | 556 | 0,98 |
| mcm4 | S | mcm4.S | NP_001079069 | ND | 0 | 0 | 0,00 | 0 | 0 | 0 | 0 | 0 | 0 | 0 | 0 |
| mcm5 | L | mcm5.L | NP_001080893.1 | TRINITY_GG_51_c364_g1_i2 | 735 | 3133 | 99,46 | 735 | 4 | 0 | 1 | 735 | 3019 | 815 | 1,00 |
| mcm5 | S | mcm5.S | NP_001080009.1 | TRINITY_GG_10_c147_g1_i2 | 735 | 3915 | 98,10 | 735 | 14 | 0 | 1 | 735 | 3834 | 1630 | 1,00 |
| mcm6 | L | mcm6.L | NP_001131039.1 | TRINITY_GG_43_c41_g1_i3 | 823 | 3231 | 98,06 | 823 | 16 | 0 | 1 | 823 | 101 | 2569 | 1,00 |
| mcm6 | S | mcm6.S | NP_001080590.1 | TRINITY_GG_70_c58_g1_i5 | 825 | 3103 | 98,18 | 825 | 12 | 1 | 1 | 825 | 70 | 2535 | 1,00 |
| mcm6.2 | L | mcm6.2.L | NP_001131039 | ND | 0 | 0 | 0,00 | 0 | 0 | 0 | 0 | 0 | 0 | 0 | 0 |
| mcm6.2 | S | mcm6.2.S | NP_001080590 | ND | 0 | 0 | 0,00 | 0 | 0 | 0 | 0 | 0 | 0 | 0 | 0 |
| mcm7 | L | mcm7.L | NP_001080722.1 | TRINITY_GG_72_c116_g1_i19 | 720 | 2579 | 98,47 | 720 | 11 | 0 | 1 | 720 | 2482 | 323 | 1,00 |
| mcm7 | S | mcm7.S | NP_001081466.1 | TRINITY_GG_11_c2882_g1_i1 | 720 | 2589 | 98,47 | 720 | 11 | 0 | 1 | 720 | 2520 | 361 | 1,00 |
| mcmbp | L | mcmbp.L | NP_001080558.1 | TRINITY_GG_37_c974_g1_i6 | 626 | 2286 | 96,01 | 627 | 23 | 2 | 1 | 626 | 150 | 2027 | 1,00 |
| mtbp | S | mtbp.S | NP_001084870 | ND | 0 | 0 | 0,00 | 0 | 0 | 0 | 0 | 0 | 0 | 0 | 0 |
| orc1 | L | orc1.L | NP_001081806.1 | TRINITY_GG_51_c930_g1_i4 | 886 | 2955 | 97,74 | 886 | 20 | 0 | 1 | 886 | 2864 | 207 | 1,00 |
| orc2 | S | orc2.S | XP_018092774.1 | TRINITY_GG_70_c1105_g1_i3 | 560 | 2172 | 97,32 | 560 | 15 | 0 | 1 | 560 | 2115 | 436 | 1,00 |
| orc3 | S | orc3.S | XP_018119715.1 | TRINITY_GG_3_c4093_g1_i1 | 709 | 2346 | 98,03 | 709 | 14 | 0 | 1 | 709 | 27 | 2153 | 1,00 |
| orc4 | L | orc4.L | NP_001081561.1 | TRINITY_GG_43_c223_g1_i6 | 432 | 1514 | 98,15 | 432 | 8 | 0 | 1 | 432 | 1381 | 86 | 1,00 |
| orc5 | L | orc5.L | NP_001089444.1 | TRINITY_GG_78_c823_g1_i6 | 448 | 1812 | 97,72 | 395 | 9 | 0 | 1 | 395 | 1808 | 624 | 0,88 |
| orc6 | L | orc6.L | NP_001086912.1 | TRINITY_GG_51_c473_g1_i6 | 225 | 1944 | 97,17 | 212 | 6 | 0 | 1 | 212 | 1585 | 950 | 0,94 |
| pcna | L | pcna.L | NP_001082364.1 | TRINITY_GG_72_c1301_g1_i1 | 261 | 1267 | 100,00 | 261 | 0 | 0 | 1 | 261 | 254 | 1036 | 1,00 |
| pcna | S | pcna.S | NP_001081011.1 | TRINITY_GG_13_c116_g1_i1 | 261 | 1158 | 99,62 | 261 | 1 | 0 | 1 | 261 | 120 | 902 | 1,00 |
| plk1 | L | plk1.L | XP_018092005.1 | TRINITY_GG_49_c568_g1_i8 | 598 | 2395 | 99,00 | 598 | 6 | 0 | 1 | 598 | 231 | 2024 | 1,00 |
| plk1 | S | plk1.S | NP_001080788.1 | TRINITY_GG_69_c53_g1_i5 | 598 | 2382 | 97,95 | 585 | 10 | 1 | 1 | 585 | 227 | 1975 | 0,98 |
| pola1 | S | pola1.S | NP_001082055.1 | TRINITY_GG_54_c103_g1_i2 | 1458 | 5058 | 97,81 | 1459 | 28 | 2 | 1 | 1458 | 4987 | 620 | 1,00 |
| pola2 | L | pola2.L | XP_018112289.1 | TRINITY_GG_50_c2973_g1_i4 | 673 | 2550 | 97,20 | 606 | 16 | 1 | 68 | 673 | 112 | 1926 | 0,90 |
| pold1 | L | pold1.L | NP_001087694.1 | TRINITY_GG_40_c131_g1_i8 | 1109 | 4117 | 98,38 | 1109 | 18 | 0 | 1 | 1109 | 4053 | 727 | 1,00 |
| pold2 | L | pold2.L | XP_018106174.1 | TRINITY_GG_72_c201_g1_i3 | 463 | 1712 | 97,84 | 463 | 10 | 0 | 1 | 463 | 1582 | 194 | 1,00 |
| pold3 | L | pold3.L | NP_001082759.1 | TRINITY_GG_87_c289_g1_i10 | 454 | 1766 | 93,83 | 454 | 27 | 1 | 1 | 454 | 116 | 1474 | 1,00 |
| pole | L | pole.L | XP_018115858 | ND | 0 | 0 | 0,00 | 0 | 0 | 0 | 0 | 0 | 0 | 0 | 0 |
| pole2 | L | pole2.L | NP_001083783.1 | TRINITY_GG_17_c1857_g1_i6 | 527 | 3215 | 99,05 | 527 | 5 | 0 | 1 | 527 | 3195 | 1615 | 1,00 |
| pole3 | L | pole3.L | XP_018084282.1 | TRINITY_GG_16_c199_g1_i6 | 147 | 1392 | 98,98 | 98 | 1 | 0 | 1 | 98 | 330 | 623 | 0,67 |
| pole4 | L | pole4.L | NP_001090193.1 | TRINITY_GG_73_c4609_g1_i1 | 116 | 860 | 96,55 | 116 | 4 | 0 | 1 | 116 | 830 | 483 | 1,00 |
| ppp1ca | L | ppp1ca.L | NP_001080222 | TRINITY_GG_48_c52_g1_i3 | 326 | 2013 | 100,00 | 326 | 0 | 0 | 1 | 326 | 171 | 1148,00 | 1,00 |
| ppp1ca | S | ppp1ca.S | XP_041434081 | TRINITY_GG_68_c39_g1_i1 | 326 | 1932 | 99,39 | 326 | 2 | 0 | 1 | 326 | 1830 | 853,00 | 1,00 |
| ppp1cb | L | ppp1cb.L | NP_001085426 | TRINITY_GG_21_c747_g1_i5 | 327 | 2590 | 100,00 | 327 | 0 | 0 | 1 | 327 | 98 | 1078,00 | 1,00 |
| ppp1cb | S | ppp1cb.S | XP_018120397 | TRINITY_GG_4_c47_g1_i4 | 327 | 1307 | 98,61 | 72 | 1 | 0 | 256 | 327 | 3 | 218,00 | 0,22 |
| ppp1cc | L | ppp1cc.L | NP_001081308 | ND | 0 | 0 | 0,00 | 0 | 0 | 0 | 0 | 0 | 0 | 0 | 0,00 |
| ppp1cc | S | ppp1cc.S | NP_001080904 | TRINITY_GG_94_c109_g2_i3 | 323 | 1931 | 100,00 | 323 | 0 | 0 | 1 | 323 | 1757 | 789,00 | 1,00 |
| recql4 | L | recql4.L | NP_001089101.2 | TRINITY_GG_86_c463_g1_i2 | 1504 | 3659 | 93,10 | 1188 | 68 | 7 | 321 | 1504 | 2 | 3535 | 0,79 |
| rfc1 | L | rfc1.L | XP_018084656.1 | TRINITY_GG_24_c253_g1_i4 | 1147 | 3514 | 96,68 | 1114 | 37 | 0 | 1 | 1114 | 3397 | 56 | 0,97 |
| rfc2 | L | rfc2.L | NP_001084837.1 | TRINITY_GG_87_c240_g1_i6 | 348 | 1414 | 99,42 | 347 | 2 | 0 | 1 | 347 | 33 | 1073 | 1,00 |
| rfc2 | S | rfc2.S | XP_041440060.1 | TRINITY_GG_53_c149_g1_i1 | 255 | 1031 | 86,46 | 229 | 30 | 1 | 1 | 228 | 988 | 302 | 0,90 |
| rfc3 | L | rfc3.L | NP_001089570.1 | TRINITY_GG_87_c1728_g1_i5 | 356 | 2471 | 99,16 | 356 | 3 | 0 | 1 | 356 | 2356 | 1289 | 1,00 |
| rfc4 | L | rfc4.L | NP_001082757.1 | TRINITY_GG_20_c735_g1_i4 | 363 | 1390 | 96,39 | 360 | 13 | 0 | 4 | 363 | 1390 | 311 | 0,99 |
| rfc5 | L | rfc5.L | NP_001080677 | ND | 0 | 0 | 0,00 | 0 | 0 | 0 | 0 | 0 | 0 | 0 | 0 |
| rfc5 | S | rfc5.S | NP_001085526.1 | TRINITY_GG_94_c389_g1_i1 | 335 | 1561 | 97,61 | 335 | 8 | 0 | 1 | 335 | 1444 | 440 | 1,00 |
| rif1 | S | rif1.S | NP_001267578.1 | TRINITY_GG_70_c107_g1_i5 | 2327 | 8327 | 93,52 | 2331 | 118 | 4 | 1 | 2327 | 136 | 7041 | 1,00 |
| rpa1 | L | rpa1.L | NP_001081585.1 | TRINITY_GG_86_c275_g1_i2 | 609 | 3567 | 99,34 | 609 | 4 | 0 | 1 | 609 | 110 | 1936 | 1,00 |
| rpa1 | S | rpa1.S | XP_041439246.1 | TRINITY_GG_54_c1677_g1_i1 | 611 | 1831 | 94,43 | 539 | 27 | 2 | 73 | 611 | 3 | 1610 | 0,88 |
| rpa2 | L | rpa2.L | NP_001085393.1 | TRINITY_GG_86_c369_g1_i1 | 276 | 1750 | 97,96 | 245 | 5 | 0 | 32 | 276 | 1528 | 794 | 0,89 |
| rpa2 | S | rpa2.S | NP_001090609.1 | TRINITY_GG_59_c378_g1_i1 | 274 | 783 | 92,13 | 216 | 15 | 1 | 28 | 243 | 140 | 781 | 0,79 |
| rpa3 | L | rpa3.L | NP_001088529.1 | TRINITY_GG_77_c471_g1_i5 | 121 | 873 | 98,31 | 118 | 2 | 0 | 1 | 118 | 71 | 424 | 0,98 |
| rpa3 | S | rpa3.S | NP_001086815.1 | TRINITY_GG_80_c1093_g1_i14 | 121 | 921 | 95,87 | 121 | 5 | 0 | 1 | 121 | 854 | 492 | 1,00 |
| ticrr | S | ticrr.S | XP_018110813.1 | TRINITY_GG_72_c316_g1_i1 | 2020 | 5764 | 83,03 | 1856 | 246 | 10 | 171 | 2020 | 5759 | 381 | 0,92 |
| topbp1 | L | topbp1.L | NP_001082568.1 | TRINITY_GG_77_c421_g1_i1 | 1513 | 5035 | 97,56 | 1513 | 37 | 0 | 1 | 1513 | 4975 | 437 | 1,00 |
| yap1 | L | yap1.L | XP_041438614.1 | TRINITY_GG_87_c1521_g1_i1 | 459 | 2402 | 98,15 | 433 | 6 | 1 | 27 | 459 | 1897 | 605 | 0,94 |
| yap1 | S | yap1.S | NP_001167495.1 | TRINITY_GG_60_c5527_g1_i3 | 335 | 1274 | 97,31 | 260 | 7 | 0 | 76 | 335 | 1228 | 449 | 0,78 |
| zmcm3 | L | zmcm3.L | NP_001080158.1 | TRINITY_GG_19_c18_g1_i5 | 806 | 3523 | 98,26 | 803 | 13 | 1 | 5 | 806 | 318 | 2726 | 1,00 |
